# Reconceptualising resilience within a translational framework is supported by unique and brain-region specific transcriptional signatures in mice

**DOI:** 10.1101/2020.11.15.383489

**Authors:** Sarah Ayash, Thomas Lingner, Soojin Ryu, Raffael Kalisch, Ulrich Schmitt, Marianne B. Müller

**Author notes:** Shared last authorship (Ulrich Schmitt, Marianne B. Müller). **Materials & Correspondence:** Correspondence should be addressed to Marianne B. Müller.

## Abstract

Chronic social defeat (CSD) in mice has been increasingly employed in experimental resilience research. Particularly, the degree of CSD-induced social avoidance is used to classify animals into resilient (socially non-avoidant) versus susceptible (avoidant). Inspired by human data pointing to threat-safety discrimination and responsiveness to extinction training of aversive memories as characteristics of resilient individuals, we here describe a translationally informed stratification which identified *three* phenotypic subgroups of mice following CSD: the *Discriminating-avoiders*, characterised by successful social threat-safety discrimination and successful extinction of social avoidance; the *Indis-criminate-avoiders*, showing aversive response generalisation, and the *Non-avoiders* (absence of social avoidance) displaying impaired conditioned learning. Furthermore, and supporting the biological validity of our approach, we uncovered subgroup-specific transcriptional signatures in classical fear conditioning and anxiety-related brain regions. Our reconceptualisation of resilience in mice refines the currently used dichotomous classification and contributes to advancing future translational approaches.

## INTRODUCTION

Stress is a well-known major risk factor for the development of mental disorders (1–2). However, the large majority of the human population does not develop stress-related mental disorders following stressor exposure (adversity/challenging life circumstances) (3). Stress research has been largely focused on the negative consequences of stressor exposure (stress susceptibility) rather than stress resilience. Focusing on stress resilience, defined as the maintenance or quick recovery of mental health during and after adversity, is a paradigm shift from disease-to health-oriented research (4). Stress resilience research has become increasingly popular over the last decade in studies on humans and experimental animals (5), the latter facilitating the search for neurobiological mechanisms underlying resilience (3; 6–10). Animal models usually employ controlled laboratory stressors to model the life stressors experienced by humans and subsequent behavioural assays, in order to determine to what extent the stressor has result in an impairment in the adaptive behaviour, relative to a non-stressed control group. Specifically, the majority of animal models are designed to investigate the role of individuality, i.e. heterogeneity across different individuals in response to the same stressor (11–12).

During recent years, chronic social defeat (CSD) has been introduced as the mouse model of choice in stress resilience research, employing social stress as a major type of adversity that also affects humans. During the CSD, male experimental mice are daily attacked for a specific number of days by another male mouse (aggressor). Every day, the aggressor mouse is a different individual but all belong to the same strain (aggressors’ strain). Subsequent behavioural characterisation employs the social interaction test (SIT), in which the experimental mouse is exposed to a novel individual mouse (social target) from the aggressors’ strain. The test allows for quantifying CSD-induced social avoidance of the novel social target in defeated compared to non-defeated control mice (13–15). Specifically, the development of social avoidance is considered a maladaptive generalised aversive response, and thus a measurement of stress susceptibility (13). In contrast, maintaining social interaction at levels comparable to non-defeated controls is considered an adaptive response, and thus a measurement of resilience (13), in analogy to the above outcome-based definition of resilience as maintained mental health after stressor exposure.

Social avoidance is a core symptom domain of social anxiety and affective disorders (16). However, avoidance behaviour can have the function to protect an individual’s well-being, and avoiding an experienced source of threat is not per se maladaptive, as much as not avoiding a threat is also not necessarily adaptive. Therefore, it remains questionable if the absence of CSD-induced social avoidance, as assessed in the SIT, is an adaptive response towards this particular stressful experience and therefore an appropriate measure of resilience in animals (15; 17–21).

Previously, we identified the involvement of conditioned learning in CSD (21). In particular, we showed that CSD-induced social avoidance is specific towards the aggressors’ strain and can be reversed following extinction training involving repeated but safe (without physical attacks) exposure to individual mice from the aggressors’ strain (21). Specifically, we designed the Modified Social Interaction Test (MSIT; 21) where defeated male mice are given the possibility to interact with a novel male individual from the aggressors’ strain and also a male individual from a novel strain, both presented simultaneously in different compartments of the test arena (21). Interestingly, defeated mice (on a group level) avoid only the former, suggesting that CSD-induced social avoidance does not generalise to mouse strains with different phenotypic characteristics but is specific to the phenotypic characteristics of the aggressors’ strain (21). Taken together, this indicates that social avoidance is an aversive conditioned response towards the aggressors’ strain, which serves as the threat-associated cue. Arguably, conditioned learning is primarily an adaptive process, protecting against threats. Therefore, CSD-induced social avoidance may also be an appropriate response towards a potential danger. In-depth analyses of the MSIT results using a larger experimental group revealed the presence of individual differences within the defeated group not only in CSD-induced social avoidance towards the aggressors’ strain but also in the way the defeated mice’ avoidance behaviour discriminated between both strains, i.e. between the threat-associated cue (aggressors’ strain) and the neutral cue (novel strain) (23).

Aiming for a behaviourally valid stratification, we hypothesised that individual differences following CSD in social avoidance development towards the aggressors’ strain and social threat-safety discrimination, as observed using the MSIT, correspond to: 1. General differences in conditioned learning assessed using an active avoidance task and 2. Responsiveness to extinction training of social aversive memories, namely CSD-induced social avoidance. Finally, to confirm the biological validity of this novel stratification, we investigated if it is reflected in unique transcriptional signatures in brain regions classically related to fear conditioning (21) and anxiety-like behaviour (24–25), specifically, medial prefrontal cortex (mPFC), ventral hippocampus (vHC), and basolateral amygdala (blA) (Figure S1. Schematic Timeline).

## RESULTS

### The MSIT identifies three distinct phenotypic subgroups within a single defeated group

During the CSD, experimental mice are attacked each day for a specific number of days by a mouse from a specific strain (aggressors’ strain; CD-1; 13), the individual aggressor changing from day to day. This design assures that the phenotypic characteristics of the aggressors’ strain, and not a particular individual mouse of the strain, become the threat-associated cue (21). Generalisation of CSD-induced social avoidance then means extension of avoidance behaviour in a later social interaction test to mice from other strains or, more specifically, to mice with other strain-associated phenotypic characteristics, while avoidance of novel individuals from the aggressors’ strain is an adaptive aversive conditioned response to the threat-associated cue (21). Threat-safety discrimination in this context would then imply to exclusively avoid mice from the aggressors’ strain. The CSD paradigm also involves a control group of non-defeated mice.

In our new paradigm, *Defeated* mice underwent the MSIT, where two novel social targets were presented simultaneously, one belonging to the aggressors’ strain, the other belonging to a novel strain with different phenotypic characteristics (21; Figure 1 and Materials and Methods). Target-specific social interaction in the MSIT is quantified as the time spent interacting with a target in the mesh enclosure relative to the time spent exploring the empty mesh enclosures (see Materials and Methods), CSD-induced social avoidance of a target being apparent from a social interaction index<1. In the current state of research using the SIT, *Resilient* and *Susceptible* subgroups within a single defeated group are identified by their social interaction index towards the single presented novel social target from the aggressors’ strain. The *Resilient* subgroup in this classical approach has an index≥1 (non-avoidant), while the *Susceptible* subgroup has a social interaction index<1 (avoidant). By contrast here, following the MSIT and calculating two social interaction indices, one for each of the novel target mice, we identified three distinct phenotypic subgroups within the *Defeated* group (Figure 1): mice with a social interaction index≥1 with the aggressors’ strain were labelled *Non-avoiders*, mice with a social interaction index<1 with both strains were labelled *Indiscriminate-avoiders*, and mice with a social interaction index≥1 exclusively with the novel strain were labelled *Discriminating-avoiders.*

**Figure 1.**
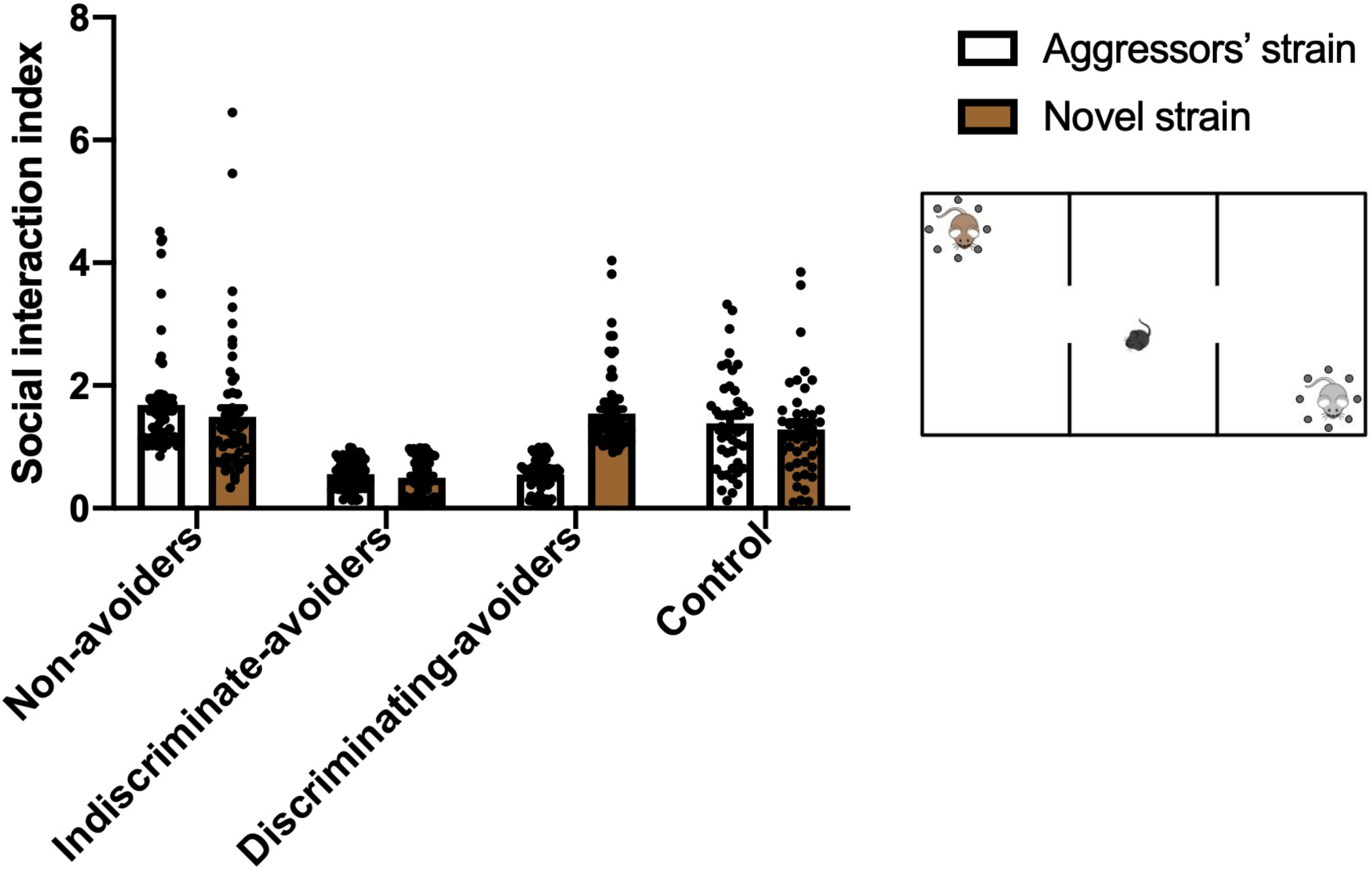
Modified Social Interaction Test (MSIT) following Chronic Social Defeat (CSD). (a) The test is performed in sociability arenas. Each arena is divided into three equal parts separated by transparent walls with openings that allow the experimental mouse (black) to move in-between. At each of the peripheries, one mesh enclosure is placed. One enclosure contains a novel mouse from the aggressors’ strain (white); the other contains a novel mouse from a novel strain matched in age, size, and sex to the novel mouse from the aggressors’ strain but of different fur colour (brown). Like the aggressor mice in the CSD, these social target mice are older and larger than the experimental mouse. The experimental mouse is introduced in the middle area of the arena and is allowed to explore the arenas for 6min. (b). The *Defeated* group was divided into three subgroups based on the animals’ social interaction indices with the social targets. Mice with a social interaction index≥1 with the aggressors’ strain were termed *Non-avoiders*, mice with a social interaction index<1 with both strains were termed *Indiscriminate-avoiders*, and mice with a social interaction index≥1 *only* with the novel strain were termed *Discriminating-avoiders.* Non-defeated *Control* mice had similar indices with both strains. Results presented as mean±s.e.m, *Non-avoiders* n=58, *Indiscriminate-avoiders* n=64, *Discriminating-avoiders* n=71, *Control* n=43.

We hypothesise the following: 1. The *Non-avoiders* subgroup’s social interaction levels with the aggressors’ strain reflects an inappropriate aversive conditioned response towards the threat-associated cue, and thus impaired aversive conditioned learning relative to the other two subgroups; 2. The *Indiscriminate-avoiders* subgroup’s social avoidance of the aggressors’ strain reflects intact conditioned learning; however, social avoidance of the novel strain reflects a generalised aversive conditioned response towards a neutral cue, i.e. failed threat-safety discrimination, rendering this subgroup the susceptible one; 3. The *Discriminating-avoiders* subgroup’s social avoidance exclusively towards the aggressors’ strain reflects intact conditioned learning, whereas social interaction levels with the novel strain comparable to the *Control* group reflect successful threat-safety discrimination, rendering this subgroup the resilient one (Table 1. Comparison between the MSIT and the SIT subgroups).

**Table 1.** Comparison between the MSIT and the SIT subgroups. The SIT presents a single novel individual mouse from the aggressors’ strain (encountered earlier during the chronic social defeat days) as a result only sociability towards the aggressors’ strain can be assessed. The development of social avoidance is considered a generalised aversive response and thus, a measurement of susceptibility. In contrast, maintaining social interaction levels comparable to non-defeated controls is a measurement of resilience. The MSIT simultaneously presents two novel individuals; one from the aggressors’ strain while the other from a novel strain, as a result sociability towards both strains can be assessed in addition to threat-safety discrimination ability. The development of social avoidance to-wards the aggressors’ strain is considered a conditioned aversive response towards a threat-associated cue whereas maintained social interaction levels comparable to non-defeated controls is the result of a failure to conditionally learn and thus, is a measurement of impaired learning. The development of social avoidance towards the novel strain is considered a generalised aversive response towards a novel and a neutral cue and thus, a measurement of susceptibility. In contrast, maintained social interaction levels comparable to non-defeated controls is a measurement of successful discrimination between a threatassociated cue (aggressors’ strain) and a neutral cue (novel strain).

| Comparison between the MSIT and the SIT subgroups |  |  |  |
| --- | --- | --- | --- |
| Test | Behaviour towards aggressors' strain | Behaviour towards novel strain | Interpretation in the context of resilience research |
| Social Interaction Test (SIT) | Interact | NA | Resilience |
|  | Avoid | NA | Susceptibility |
| Modified Social Interaction Test (MSIT) | Interact | - | Impaired learning |
|  | Avoid | Avoid | Susceptibility |
|  | Avoid | Interact | Resilience |

### Impaired ability to conditionally learn in the Non-avoiders subgroup in a multi-day operational learning task

If the results in the MSIT reflect true ability, or lack thereof, to conditionally learn an aversive social cue, then these individual behavioural differences should extend to aversive non-social cues (hypothesis 1-3). Following CSD and MSIT, a first subset of animals from each of the three *Defeated* subgroups underwent an operational learning task in which animals had to acquire active avoidance behaviour over multiple days (Figure S1). The task involved two dimensions. The first is learning that the tone signals the upcoming electrical foot shock, and the second is to employ the appropriate fear reaction of this learned association by fleeing to the other side of the arena, instead of freezing (fright). All subgroups were able to learn the task, indicated by a significant increase in conditioned (correct) response% throughout the training days as revealed by two-way ANOVA where days (the factor with repeated measures) was significant (F (5, 160)=46.72, p<0.0001***; Figure 2). However, the *Non-avoiders* subgroup did so to a lesser extent as revealed by a significant effect of the subgroups factor and the interaction between the subgroups and time factors (Subgroups F (2, 32)=4.359, p=0.0212*, Interaction F (10, 160)=3.042, p=0.0015**; Figure 2). Moreover, the *Non-avoiders* subgroup had a significantly lower increase in conditioned response% on the seventh day compared to the *Indiscriminate-avoiders* and the *Discriminating-avoiders* subgroups (Bonferroni, p<0.01**; Figure 2). Furthermore, the *Non-avoiders* subgroup was the only one not to reach a conditioned response% above chance level (50%) on the last day whereas the *Discriminating-avoiders* and *Indiscriminate-avoiders* subgroups were significantly above chance level (One sample t-test, *Non-avoiders* (t=0.5490, df=17, p=0.5902), *Discriminating-avoiders* (t=2.97, df=8, p=0.0178*), *Indiscriminate-avoiders* (t=3.651, df=9, p=0.0053**); data not shown). Finally, conditioned response% in the last day and social interaction indices of the *Defeated* group with the aggressors’ strain in the MSIT showed a significant negative correlation (Pearson correlation, r=-0.3304, p=0.0491*, n=37; data not shown). The results suggest that social avoidance of the aggressors’ strain reflects intact learning of threat-associated cues. In contrast, maintaining normal levels of interaction with the aggressors’ strain, as seen in the *Non-avoiders* subgroup, is a reflection of impaired conditioned learning, supporting hypotheses 1-3.

**Figure 2.**
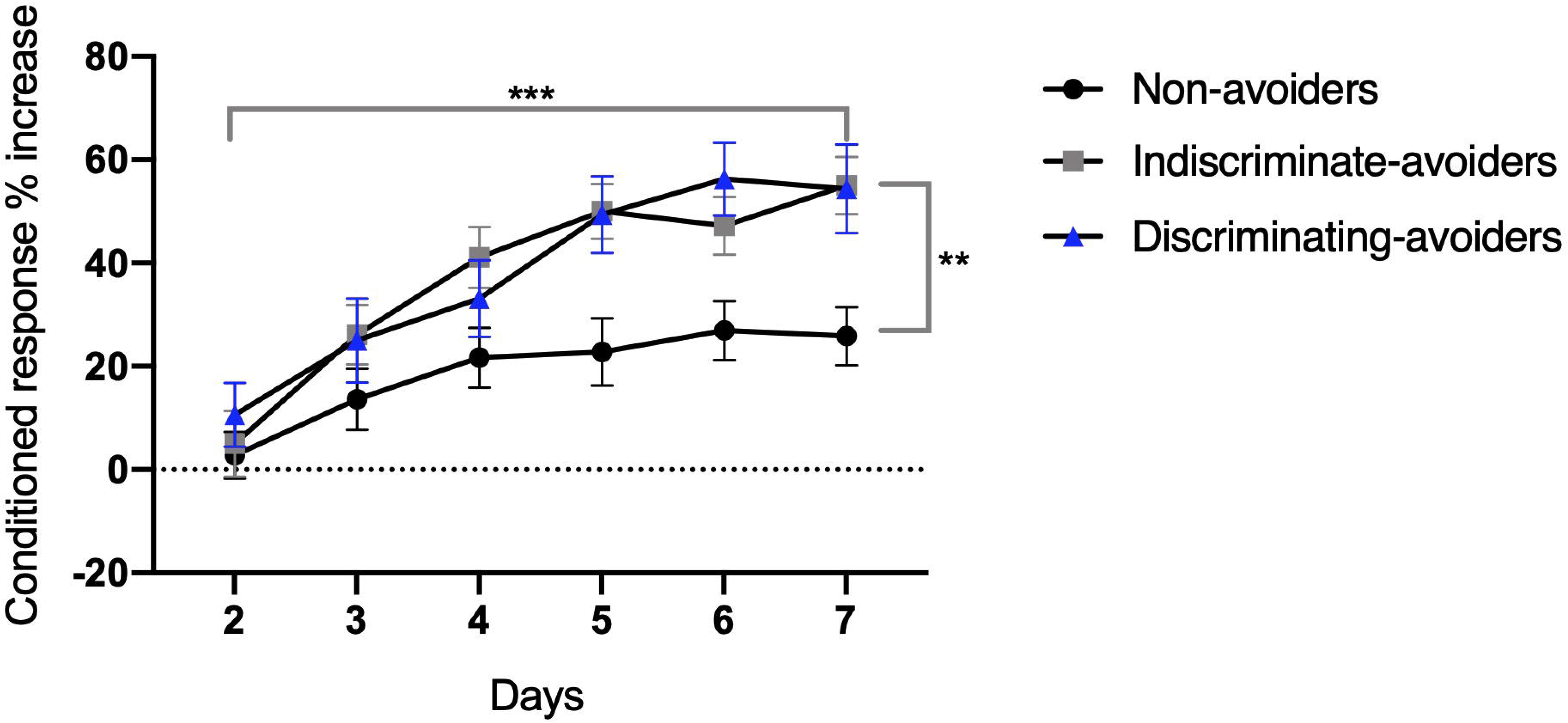
Active Avoidance Task. All subgroups were able to learn the task, indicated by a significant increase in conditioned (correct) response% throughout the training days. However, the *Non-avoiders* subgroup had a significantly lower increase in conditioned response% on the seventh day compared to the *Indiscriminate-avoiders* and the *Discriminating-avoiders* subgroups. Results presented as mean±s.e.m, two-way ANOVA, Days (factor with repeated measures): F(5, 160)=46.27, p<0.0001***, Subgroups: F(2, 32)=4.359, p=0.0212*, Interaction: F(10, 160)=3.042, p=0.0015**, and Bonferroni’s multiple comparisons test, p<0.01**, *Non-avoiders* n=18, *Indiscriminate-avoiders* n=10, *Discriminating-avoiders* n=9.

### Resistance to extinction training of aversive memories in the Indiscriminate-avoiders subgroup

Human data point to threat-safety discrimination and responsiveness to extinction training of aversive memories as characterising resilient individuals (22). Moreover, Krishnan and colleagues suggest that preserved extinction learning ability might help reduce CSD-induced generalisation of social avoidance (17). Previously, we successfully reversed CSD-induced social avoidance by applying a social avoidance extinction treatment (21). We therefore here hypothesised that, among the two *Defeated* subgroups that acquire social avoidance behaviour during CSD (*Indiscriminate-avoiders* and *Discriminating-avoiders*), only the subgroup with successful threat-safety discrimination, i.e. the resilient subgroup (*Discriminating-avoiders*) would respond to extinction (hypotheses 2-3). To this end, following the MSIT, a second subset of animals from each of the two *Defeated* sub-groups underwent social avoidance extinction treatment followed by a second MSIT (Figure S1). The treatment involved repeated visual and olfactory exposure to the same aggressors encountered during the CSD without any physical attack, as a result of maintaining a mesh wall between the animals (see Materials and Methods). Using two-way ANOVA, where the extinction treatment (the factor with repeated measures) and the MSIT subgroups were considered as two factors, significant effects were found for both factors and for the interaction between them (Extinction treatment: F(1, 41)=7.092, p=0.0110**, Subgroups: F(1, 41)=7.061, p=0.0112**, Interaction: F(1, 41)=8.293, p=0.0063**). Bonferroni test on the social interaction indices with the aggressors’ strain between the two MSITs (after CSD treatment and after extinction treatment, respectively) revealed that the *Discrimi-nating-avoiders* subgroup had a significantly greater index following the extinction treatment compared to before the treatment (following CSD treatment only; p=0.0002***; Figure 3). In contrast, the *Indiscriminate-avoiders* subgroup maintained similar indices (p>0.9999; Figure 3). The results reveal that both subgroups similarly socially avoided the aggressors’ strain following CSD; however, *only* the *Discriminating-avoiders* subgroup successfully extinguished their social avoidance following the extinction treatment, whereas the *Indiscriminate-avoiders* subgroup was resistant to the treatment. This supports hypotheses 2-3.

**Figure 3.**
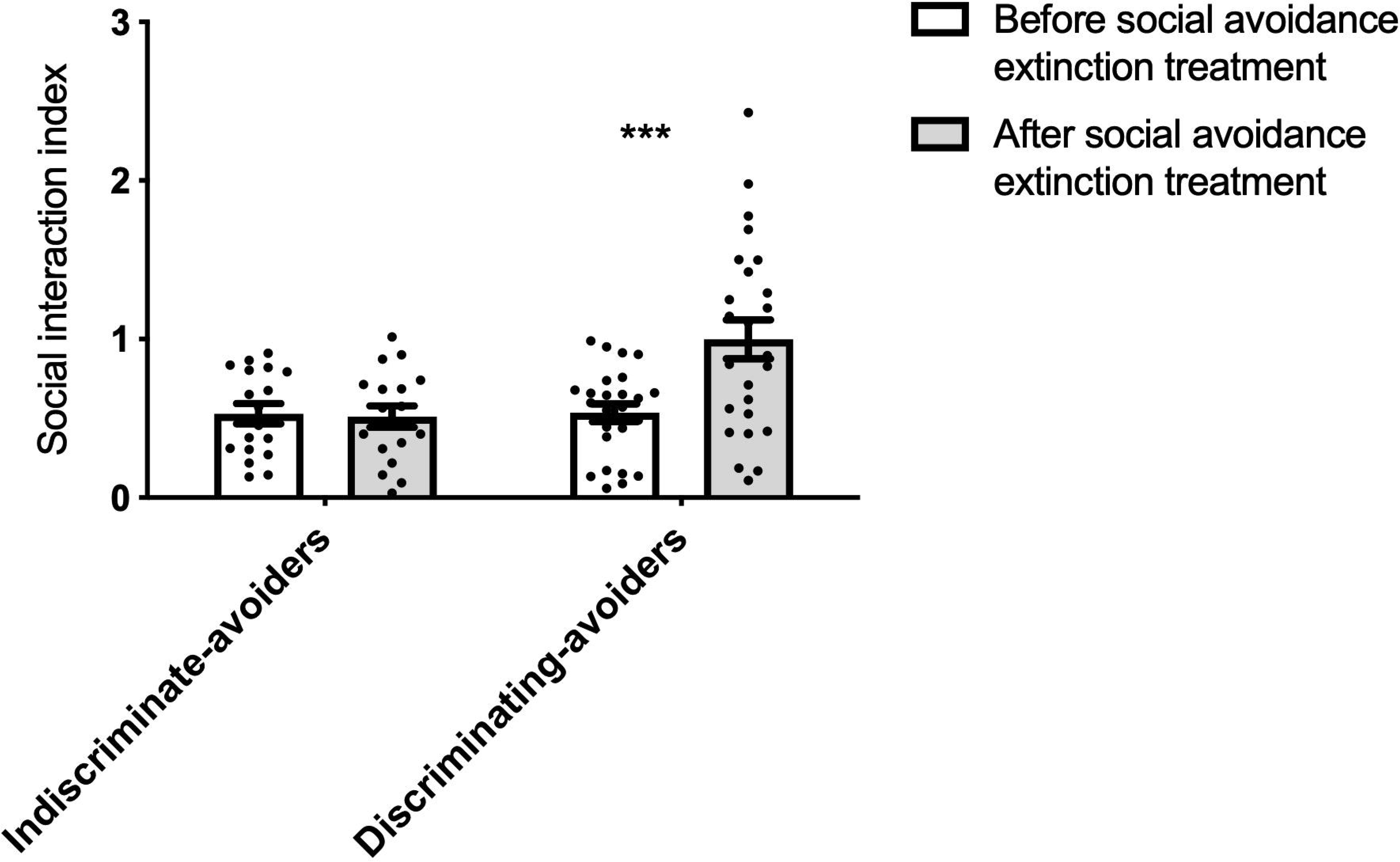
Modified Social Interaction Test following Social Avoidance Extinction Treatment. The *Discriminating-avoiders* subgroup had significantly greater social interaction index with the novel animal from the aggressors’ strain following Social Avoidance Extinction Treatment compared to their index before the treatment (following chronic social defeat only), whereas the *Indiscriminate-avoiders* subgroup maintained similar indices between the two time points. Results presented as mean±s.e.m, two-way ANOVA, Extinction treatment (factor with repeated measures): F(1, 41)=7.092, p=0.0110**, Subgroups: F(1, 41)=7.061, p=0.0112**, Interaction: F(1, 41)=8.293, p=0.0063**, and Bonferroni’s multiple comparisons test, p=0.0002***, *Indiscriminate-avoiders* n=21 and *Discriminating-avoiders* n=25.

### *Stratification using the MSIT is supported by unique* and *brain region-specific transcriptional signatures*

To confirm the biological validity of the behavioural stratification achieved using the MSIT, we performed transcriptome analysis immediately following the MSIT in three brain regions: mPFC, vHC, and, blA (Figure S1). The three regions are part of a brain circuit implicated in conditioned learning, extinction learning, and generalisation of fear (aversive) responses (23). Additionally, the same regions were described as neuroanatomical correlates of stress-related psychopathologies, including alterations in social interaction behaviour (24–25). In each region, we profiled differential gene expression between each of the three *Defeated* subgroups in comparison to the *Control* group.

Heat maps of log2 fold change of the DEGs in each brain region shows unique transcriptional signatures of each subgroup (Figure 4a). The total number of differentially expressed genes (DEGs) shared between all of the three *Defeated* subgroups was remarkably small, ranging from zero in the vHC to 12 in the blA (Figure 4b). Moreover, the number of shared genes between any two subgroups is considerably smaller than the numbers of genes differentially regulated in only one group (Figure 4b). Summing across regions, the lowest total number of DEGs (52 genes; Table S1) was found in the *Discriminating-avoiders* subgroup whereas the greatest number of DEGs was found in the *Indiscriminate-avoiders* subgroup (333 genes, Table S1), particularly in the blA (Figure 4b; Figure S2a. Volcano Plot). The *Discriminating-avoiders* subgroup was characterised by unique transcriptome signatures in all three brain regions. In fact, this subgroup was the only one to have differentially down-regulated genes in the vHC (Figure 4b; Figure S2b. Volcano Plot). On the other hand, the *Non-avoiders* subgroup was characterised by a large number of DEGs in the mPFC (Figure 4b; Figure S2c. Volcano Plot; Table S1) but no significant enrichment of any biological processes was detected in this region. However, targeted enrichment analysis of the mammalian target of rapamycin (mTOR) pathway confirmed its involvement in the mPFC in the differentially up-regulated genes of *Non-avoiders* subgroup (Enrichr online tool, p-value=0.0548*, data not shown).

**Figure 4.**
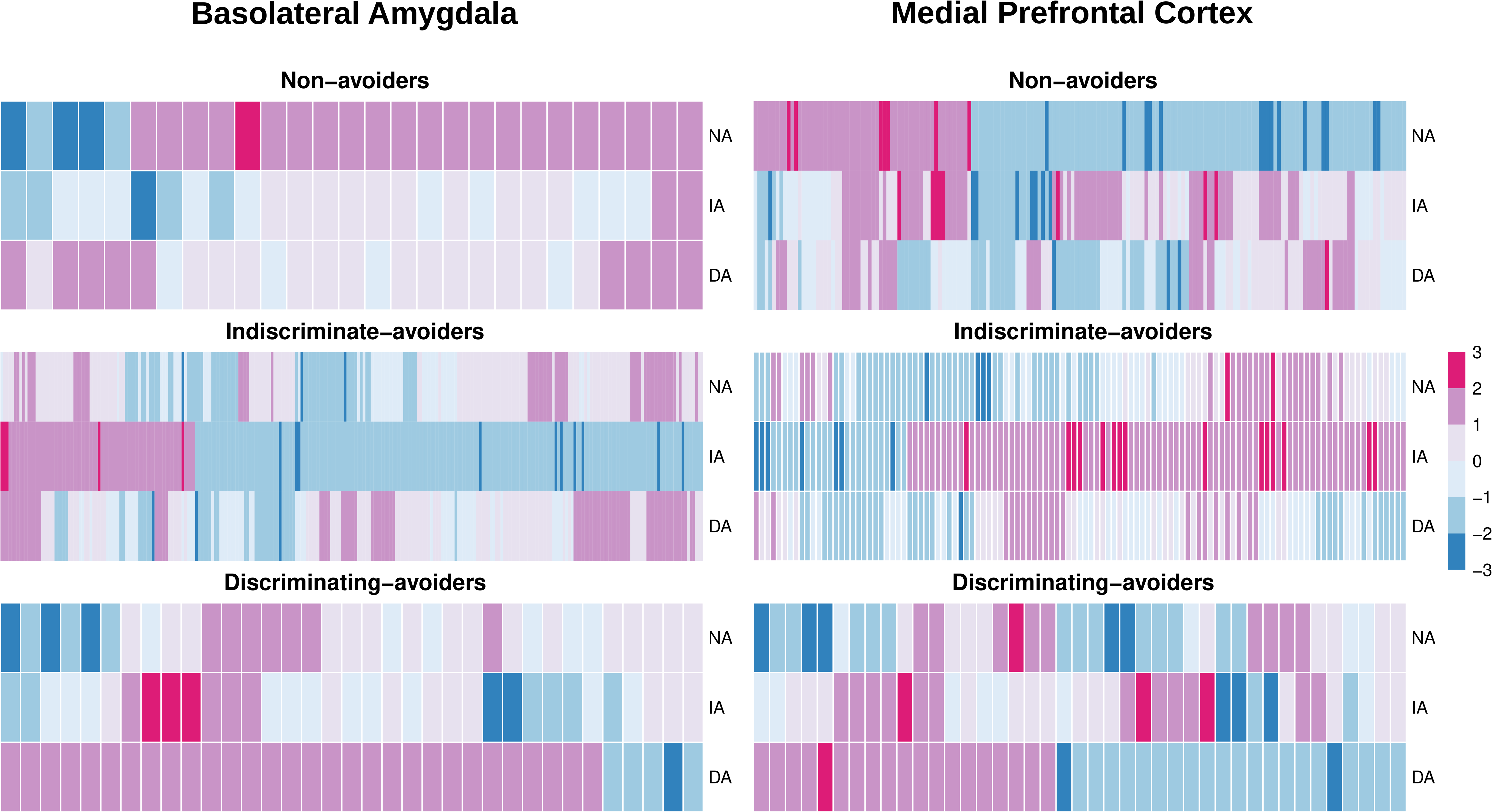
Transcriptome Analysis after Chronic Social Defeat. (a) Heat maps illustrating the regulation of genes for each condition and anatomical structure, with magenta to blue gradient depicting an up-to down-regulation (≥2-fold increase to ≤2-fold decrease). Each heat map depicts the log2-fold-change values from all comparisons for genes that were identified as differentially regulated in a particular comparison. (b) Venn diagram show the number of differentially expressed genes in the *Non-avoiders*, *Indiscriminate-avoiders*, and *Discriminating-avoiders* subgroups each compared to non-defeated controls, with the overlap depicting genes that were identically differentially expressed by the subgroups. Up-regulated (magenta) and down-regulated (blue) genes are shown separately (criteria for significance: ≥2-fold change compared to the respective anatomical Control group at p≤0.05*). NA: *Non-avoiders*, IA: *Indiscriminate-avoiders*, and DA: *Discriminating-avoiders*. n=7 per subgroup.

Enrichment analysis of the down-regulated DEGs in the blA of the *Indiscriminateavoiders* subgroup revealed suppression of biological processes involved in neuronal plasticity and learning, such as, among others, axon development, nervous system development, and regulation of axonogenesis (Figure 5a). At the phenotypic level, gene clusters like synaptic vesicle number, dendrite spine morphology, and central nervous system synaptic transmission were significantly enriched (Figure 5b). In fact, some of those downregulated genes modulate neuronal actin dynamics and synaptic plasticity (Table S1). Identifying transcription factors potentially responsible for observed changes in gene expression is necessary for understanding gene regulatory networks. Analysis of upstream regulators revealed substantial overlap between the differentially down-regulated genes in the blA and three transcription factors, specifically, J*arid2*, *Suz12*, and *Rest (*Figure 5c). All three transcription factors play crucial roles in modulating different functional aspects of the central nervous system.

**Figure 5.**
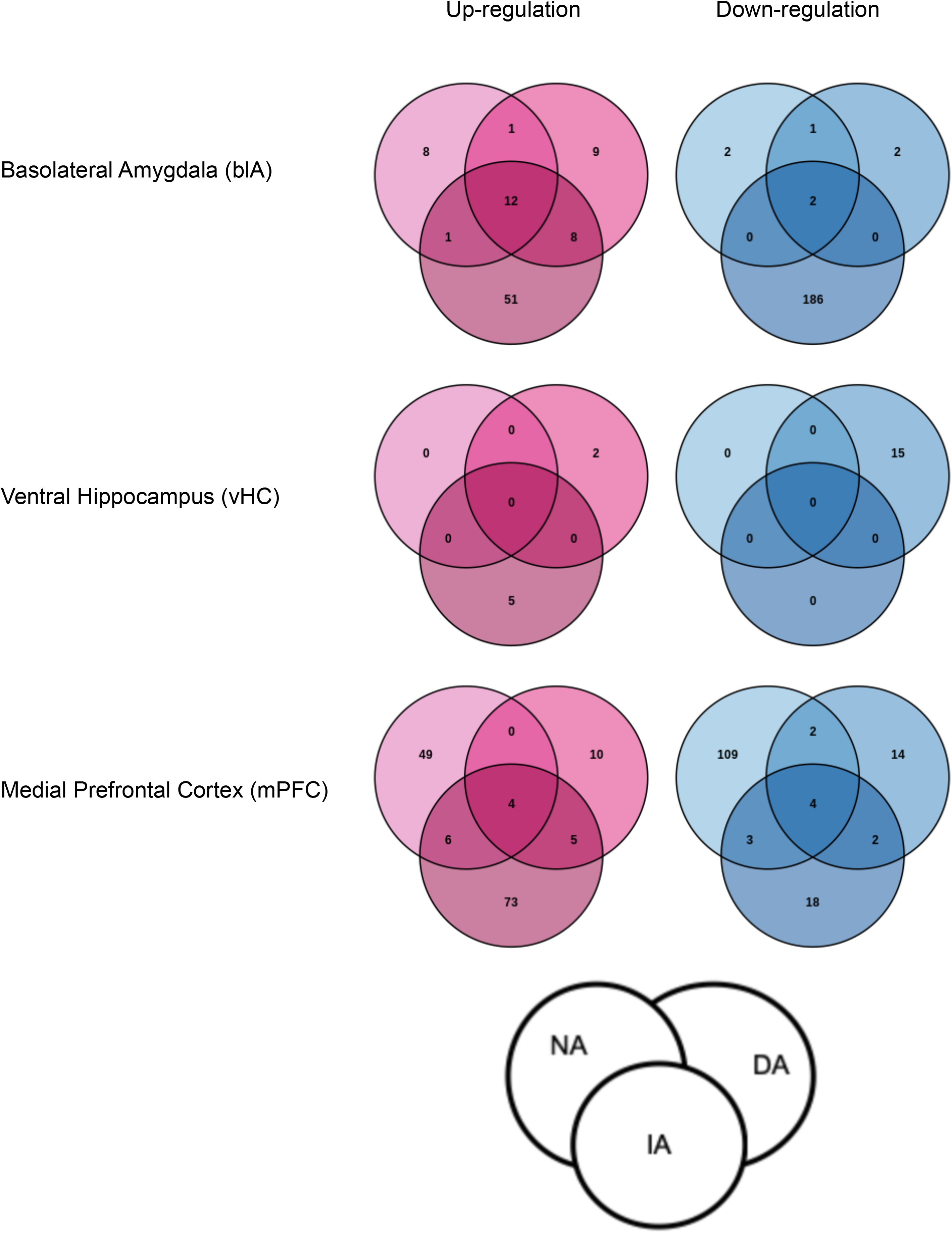
Enrichment Analysis of the differentially expressed genes in the basolateral Amygdala of the *Indiscriminate-avoiders* subgroup compared to the *Control* group. (a) Biological processes involved are mainly of axon and nervous system development, (b) Phenotypical abnormalities mainly found in synaptic vesicle number, (c) Transcription factors that substantially overlapped with the candidate genes. *Control* n=5 and *Indiscriminate-avoiders* n=6.

**Figure 6.**
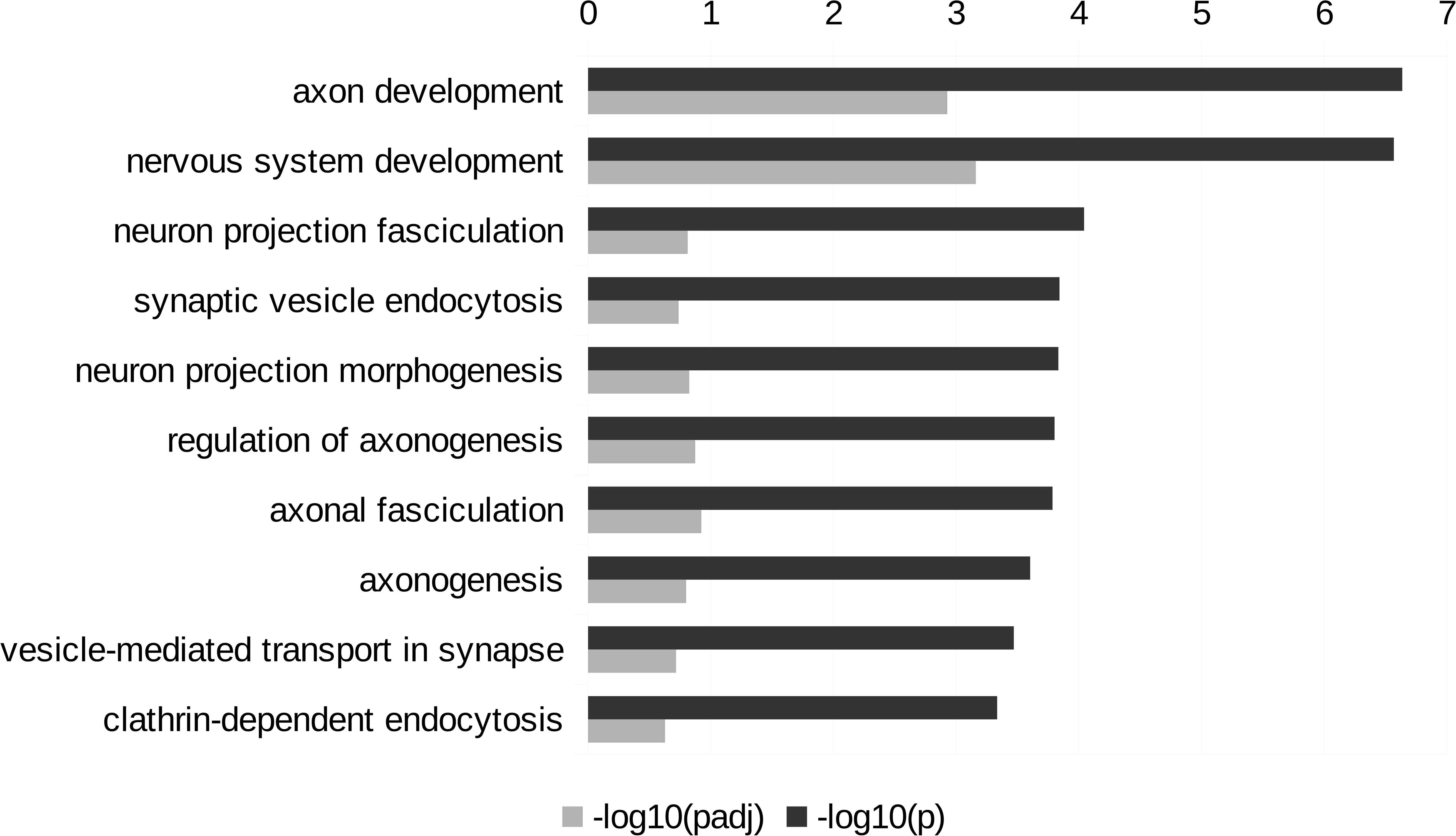
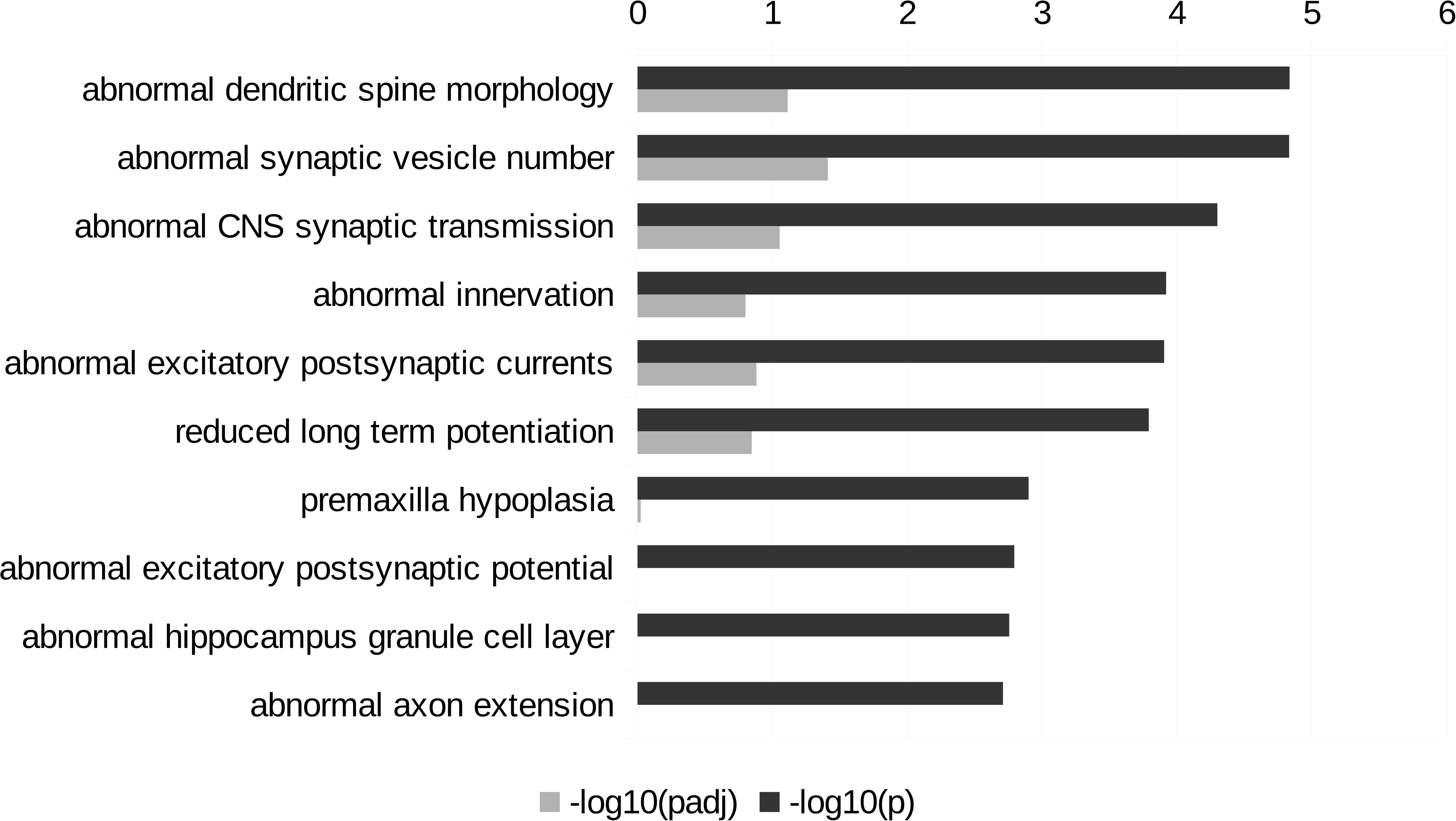
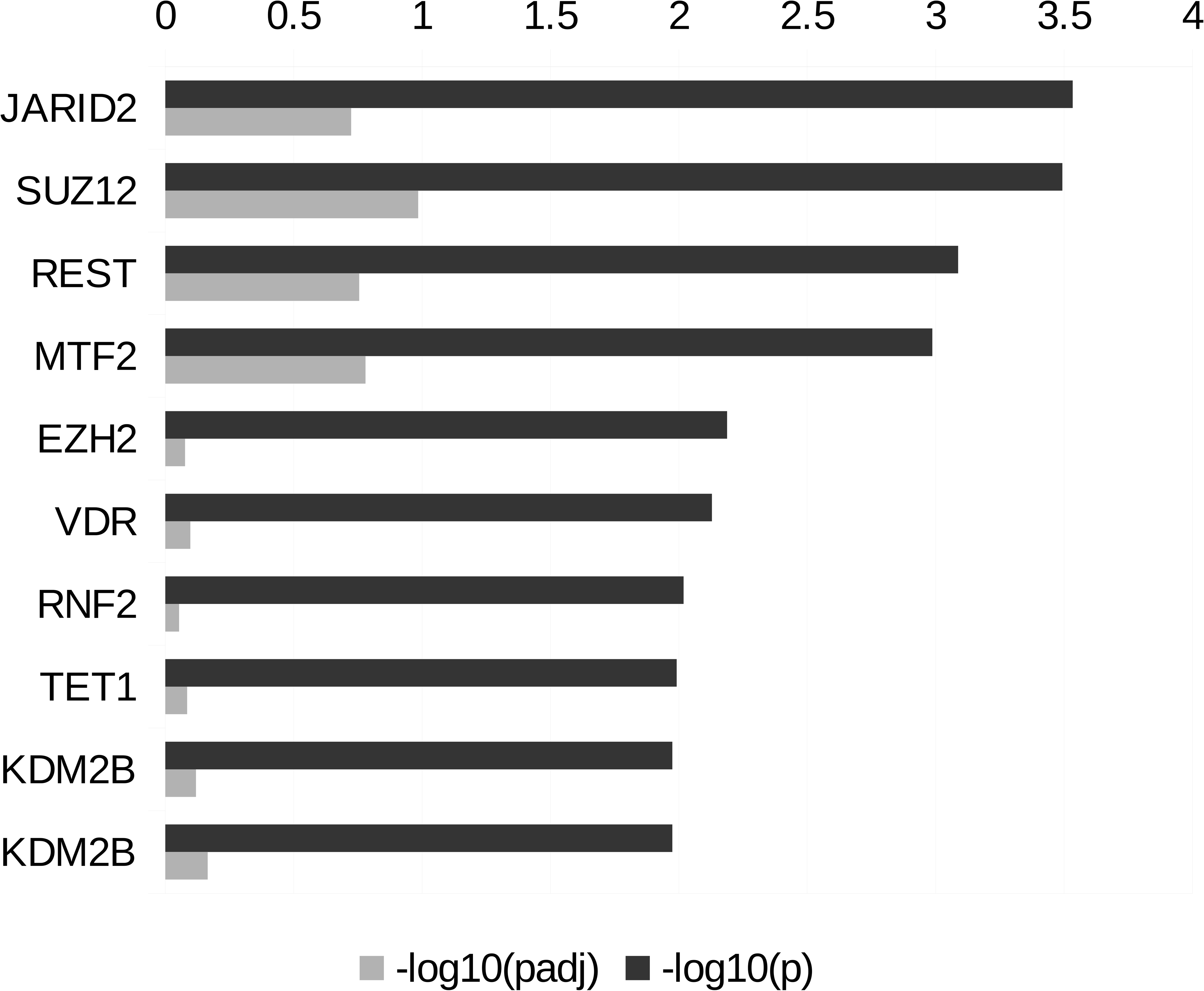
Conceptual Graphical Overview. Following chronic social defeat and using the Modified Social Interaction Test, the single group of defeated animals is stratified into three subgroups based on threat-safety discrimination and sociability. The *Discriminating-avoiders* subgroup is characterised with successful threat-safety discrimination and responsiveness to extinction training of aversive memories both of which have been discussed as characteristics of resilient individuals after the occurrence of stressful events. In contrast, the *Indiscriminate-avoiders* subgroup is characterised with aversive/fear response generalisation to similar but novel/neutral stimuli and resistance to extinction training of aversive memories both of which are key symptoms of stress-related mental disorders. Finally, the *Non-avoiders* subgroup is characterised by an impaired ability to conditionally learn.

Taken together, our results show that the novel behavioural stratification into three subgroups using the MSIT following CSD, is supported by unique and brain regions-specific transcriptional signatures in classical fear conditioning-and anxiety-related brain regions indicating that these subgroups are also different at the biological level.

## DISCUSSION

Animal experimental research into mechanisms of resilience has been largely built on a single phenotypic readout, specifically, CSD-induced social avoidance towards a novel mouse from the aggressors’ strain as assessed during the SIT (13–15). The underlying hypothesis is that social avoidance after CSD reflects a generalised aversive response (13–15). Given the continuously increasing number of publications using this approach to stratify stressed animals into R*esilient* (non-avoidant) and S*usceptible* (avoidant) subgroups, we have identified the need to: 1. Dissect the neurobehavioral mechanisms underlying the social avoidance phenotype and 2. Develop a refined and translationally informed conceptual framework to advance resilience research.

In contrast to the idea of a generalised (unspecific) aversive response, our recent data show that, on a group level, CSD-induced social avoidance is a conditioned response that is specific to the phenotypic characteristics of the aggressors’ strain (21). The simultaneous presentation of two phenotypically distinct mouse strains in the MSIT allows the assessment of two phenotypic readouts, namely CSD-induced social avoidance development and threat-safety discrimination. In-depth behavioural phenotyping using the MSIT identified three subgroups within the *Defeated* group. The *Discriminating-avoiders* subgroup developed conditioned social avoidance only towards the aggressors’ strain, without generalising their avoidance response to the novel strain, reflecting an intact aversive conditioned learning and successful threat-safety discrimination. Further, *Discriminating-avoiders* extinguished their avoidance response towards the aggressors’ strain following social avoidance extinction training. In contrast, the *Indiscriminate-avoiders* subgroup developed conditioned social avoidance towards the aggressors’ strain, reflecting an intact aversive conditioned learning however their avoidance response was generalised to the novel strain reflecting a failure in threat-safety discrimination. *Indiscriminate-avoiders* also failed to extinguish their avoidance response towards the aggressors’ strain. Lastly, the *Non-avoiders* subgroup showed continuous interaction with the aggressors’ strain, reflecting an impaired aversive conditioned learning. This was further supported by the same subgroup’s poor conditioned response in the active avoidance task and the negative correlation between this response and social interaction levels with the aggressors’ strain during the MSIT.

Generating appropriate defensive behaviours when facing threat-associated stimuli is essential for survival while withholding those behaviours when facing novel and neutral stimuli is adaptive. The characteristics of the *Discriminating-avoiders* subgroup, i. e. successful learning of threat-associated cues, threat-safety discrimination, responsiveness to extinction learning of negative memories, and mental flexibility are all discussed as characteristics of stress resilient individuals (22; 26–30). In contrast, generalised aversive/fear responses and resistance to extinction learning after the occurrence of traumatic events, as seen by the *Indiscriminate-avoiders* subgroup, are key symptoms of stress-related mental disorders such as post-traumatic stress disorder and anxiety disorders (22; 31–34) and thus, are characteristic of stress susceptible individuals. Our detailed behavioural characterisation strongly supports the validity of the MSIT as a re-classification test of resilience phenotypes in mice following CSD. (Figure 7. Conceptual Graphical Overview).

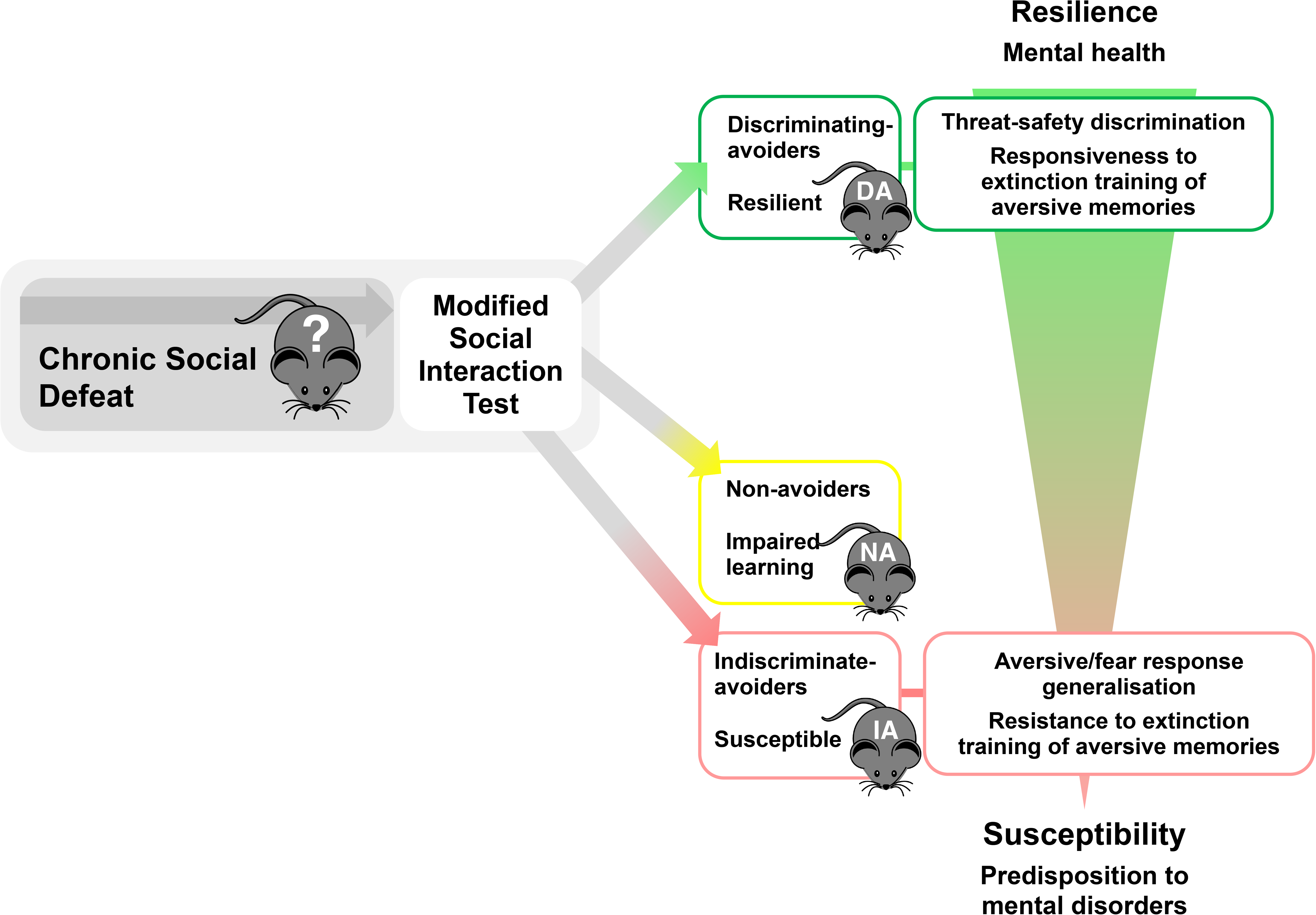

The biological validity of the MSIT’s re-classification was confirmed by the presence of distinct and biologically meaningful transcriptional signatures in classical fear conditioning-and anxiety-related brain regions, namely, the mPFC, vHC, and blA. Specifically, the blA is involved in the acquisition, prevention, and attenuation of extinction (23; 35). A considerable number of genes associated with synaptic plasticity were differentially down-regulated in the blA of the *Indiscriminate-avoiders* subgroup which showed generalised aversive/fear responses and resistance to extinction treatment. Among those plasticity-associated genes we found e.g. Synaptopodin (*synpo*; 36), the parkinson’s disease-associated synaptic vesicle endocytosis protein *dnajc6/auxilin* (37), *Neurotrophin 5* (*ntf5;* 38), glutamate ionotropic receptor kainate type subunit 5 (*grik5; 39*), the autism spectrum disorder-associated candidate *shank2* (40), and the transcription factor *neurod1* which plays a crucial role in neuronal cell fate specification (41). Plasticity adaptations are crucially involved in adaptive learning and memory processes including emotional and social memories (42). Moreover, these adaptations enable neuronal networks to stabilise activity in the face of external perturbations. The unique transcriptional signatures in the blA of the *Indiscrimi-nate-avoiders* subgroup might suggest that those animals lack the ability to flexibly adapt and consequentially fail to successfully respond to the repeated stressors they experience during the CSD. This is further supported by the substantial overlap found between the differentially down-regulated genes in the blA of the *Indiscriminate-avoiders* subgroup and the three transcription factors J*arid2*, S*uz12*, and R*est*. All three transcription factors play crucial roles in modulating different functional aspects of the central nervous system e.g. homeostatic plasticity adaptations, (R*est*; 43), neuronal activity (J*arid2;* 44), and myelination *(Suz12;* 45). In particular, *Rest* is essential for the experience-dependent fine-tuning of gene expression involved in synaptic activity and plasticity and is disturbed by early life stress (46).

The differential down-regulation of the autism spectrum disorder candidate gene *shank2* is of particular interest in the context of social behaviour being conceptualised as a core symptom of mental disorders across different diagnostic categories (47). The MSIT allows to investigate social affiliative behaviour while integrating individual information on conditioned learning and threat-safety discrimination abilities following the CSD experience. In fact, the MSIT closely resembles the three-chambered sociability task frequently used to assess autism-related phenotypes of social impairment. Shank proteins (Shank 1, −2 and −3) are multi-domain scaffolding proteins and signalling adaptors enriched at excitatory neuronal synapses, interacting with a multitude of synaptic proteins. Mutations in the human *shank2* gene have been associated with autism spectrum disorder and intellectual disability (48). Surprisingly, little information is available on a putative role of Shank proteins in modulating stress-related neurobehavioral outcomes (49). To the best of our knowledge, this is the first report pointing to a specific involvement of *shank2* in modulating *social* stress-related phenotypes.

The vHC is a critical region for memory encoding and maintaining the specificity of memories (50). Recently, it has been shown that small lesions in the vHC led to over-generalisation of the conditioned freezing response to a neutral context in rats (51). The *Discriminating-avoiders* subgroup, which is characterised by successful threat-safety discrimination, was the only subgroup displaying a substantial number of differentially down-regulated genes in the vHC. Here, the most strongly down-regulated candidate gene was angiotensin converting enzyme (*ace*; see Table S1). The renin-angiotensin-system and angiotensin converting enzyme in particular, have been repeatedly discussed as susceptibility factors for depression and the bi-directional relationship between depression and car-diovascular disorders (52–54). In accordance with our results suggesting that cognitive performance is essential to successfully cope with aversive situations and in line with the differential down-regulation of *ace* in the *Discriminating-avoiders* subgroup, studies in transgenic mice have shown that continuous activation of the central renin-angiotensin-system impairs cognitive function in an active avoidance task (55).

Finally, the *Non-avoiders* subgroup that reflected a phenotype of impaired aversive conditioned learning, had the greatest differential expression in the mPFC, a key region for various forms of adaptive learning (56). Targeted enrichment analysis of the mTOR pathway confirmed its involvement in the mPFC in the differentially up-regulated genes of this subgroup. Impairments in social behaviour in mice has been shown to be associated with mTOR pathway’s over-activation in the mPFC (57).

A limitation of our research includes treating the mPFC as a homogenous brain region and not taking into consideration its sub-anatomical areas in the transcriptome analysis. This might explain why we could not detect significant enrichment of any biological processes in this brain region. Similarly, the dorsal HC was not included in our transcriptome analysis, but only the vHC.

### Implications and future directions

To answer the question “what can be considered a meaningful phenotype of resilience in the mouse” and to strengthen the translational power, we first have to understand the neurobehavioral mechanisms shaping the phenotypic readouts observed following stress exposure. Inspired by evidence from human data pointing to threat-safety discrimination and responsiveness to extinction training of aversive memories as characteristics of resilient individuals, we have identified individual abilities shaping the development of *three* distinct behavioural subgroups within a single defeated group of mice. Our approach substantially refines the currently used dichotomous behavioural classification into *resilient* and *susceptible* individuals. Furthermore, our novel behavioural stratification is supported by unique and subgroup specific transcriptional signatures in key brain regions involved in classical fear conditioning and related to anxiety. While further studies are needed to extend our analyses to female mice to decode potential sex-specific differences in the development of stress-related mental disorders, we are confident that our proposed re-conceptualisation of resilient phenotypes in mice can contribute to substantially advance future translational approaches in resilience research.

## Supporting information

Table S1 Basolateral Amygdala

Table S1 Medial Prefrontal Cortex

Table S1 Ventral Hippocampus

## Acknowledgements

This work was supported by the Deutsche Forschungsgemeinschaft (DFG, CRC 1193 subprojects B01, C01, Z02) and the Boehringer Ingelheim Foundation. We would like to thank Dr. Konstantin Radyushkin, Dr. Inge Sillaber, Ms. Sandra Reichel, Mr. Niklas Schmitz, Dr. Anna Carboncino, Ms. Julia Deuster, Ms. Jennifer Klüpfel, and Ms. Verena Opitz, for their technical assistance. In addition, we would like to thank Dr. Maria Mendez-Lago and Ms. Hannah Lukas from the Genomics Core Facility at the Institute of Molecular Biology (Mainz, Germany) for their technical support with next generation sequencing. The funding sources had no involvement in the study design, in the collection, analysis, and interpretation of data; in the writing of the report, and in the decision to submit the article for publication.

## Author Contributions

S.A. conceived the idea, S.A., M.M., and U.S. developed the theory, S.A. planned and carried out the experiments, S.A. performed the behavioural analysis, T.L. performed the bioinformatic analyses, S.A. and S.R. supported the bioinformatic analysis, M.M., U.S., and R.K. verified the analytical methods, S.A., M.M., and U.S. interpreted the results, R.K. supported the results interpretation, M.M. supervised the project, U.S. helped supervise the project, S.A. wrote the manuscript with support from M.M., U.S., and R.K. All authors contributed to the writing of the final manuscript.

## Competing Interests

The authors declare that they have no conflict of interest. Raffael Kalisch receives advisory honoraria from JoyVentures.

## MATERIALS AND METHODS

### Animals

C57BL/6J male mice (n=236) weighing between 22-28g at the age of seven weeks were obtained from Janvier (France). All mice were housed individually and maintained in a temperature-and humidity-controlled facility on a 12hr light-dark cycle (lights on 8:00; lights off: 20:00; 23°C; 38% humidity) with food and water ad libitum. Mice were habituated for one week to the facility after arrival and before the beginning of the experiment. Tests were conducted during animals’ light phase (8:30-13:30). Procedures were performed in accordance with the European Communities Council Directive and approved by local authorities (Landesuntersuchungsamt Rheinland-Pfalz). Figure S1. Schematic Timeline.

### Chronic Social Defeat (CSD)

The treatment was performed similar to Golden et al., 2011 (13). In brief, every day for 10 straight days, mice (*Defeated* n=193) were introduced to a different home cage of a larger, older, and retired male breeder from the CD-1 strain (aggressors’ strain, pre-existing in the facility). After being physically defeated (physical phase), a mesh wall was introduced between the two mice allowing sensory contact but not physical contact (sensory phase) overnight. Age-matched mice maintained in the same conditions but randomised to the non-defeated control group (*Control* n=43) were placed for 90s in an empty cage before being returned to their individual cages, which were separated in half by identical mesh walls used for the *Defeated*. This was conducted every day throughout the 10 days. All cages were maintained in environmentally controlled cabinets (Uniprotect NG by Zoonlab) separated for each group. On the last day of the CSD procedure, all mice were housed individually in new cages and left to rest overnight. Light conditions in all tests were 37lx.

### Modified Social Interaction Test (MSIT)

The test was performed similar to Ayash et al., 2020 (21). In brief, mice were introduced in the middle area of the three-chambered arenas (total size: 60×40cm) with empty mesh enclosures, one at each of the two peripheries, to explore as the habituation phase. Following immediately from habituation, two mice from different strains were presented simultaneously each in an enclosure. One enclosure contained a novel CD-1 mouse (aggressors’ strain); the other contained a novel mouse of matching size, age, and sex to the CD-1 mouse, i.e. the mouse was a larger and older male than the mice of interest, but of different fur colour (brown) from the 129/Sv strain (novel strain, pre-existing in the facility).

Both phases lasted 6min each. The interaction zone was defined as 2cm around the mesh enclosures’ boundaries. Time spent interacting with each of the presented mice (when the nose was within the interaction zone) was scored as a measurement of social interaction levels. The social interaction index with the each of the two presented strains was calculated as follows: (time spent exploring each strain during the testing phase / average of time spent exploring the two empty mesh enclosures during the habituation phase) × 100.

### Active Avoidance Task

The test was performed in Active Avoidance boxes by TSE, Bad Nauheim, Germany. The arenas were rectangle, divided into two equal parts by smooth black walls with a small opening in between, has black walls, and metal grid floors (total size: 30×12×25cm). Animals underwent 60s of habitation followed by 10kHz tone, then depending on each animal’s behaviour the following sequence took place:

1. For animals that did not change sides within 5s, the tone was switched off and a foot shock of 0.4mA was administered until sides were changed. However, for those animals that still did not change sides, the foot shock went off after 10s duration followed by intratest-interval of 30s.
2. For animals that did change sides, the tone was played and no foot shock was applied; 30s of intra-test-interval took place immediately.

This cycle repeated itself 20× for each animal per day before 60s of delay took place. The sessions lasted seven straight days and time taken to change side during every cycle was scored as a measurement of a conditioned response.

### Social Avoidance Extinction Treatment

The test was performed similar to Ayash et al., 2020 (21). In brief, *Indiscriminate-avoiders* and *Discriminating-avoiders* subgroups, as identified using the MSIT, alternated between the same cages of the previously encountered aggressors during the CSD. However, a mesh wall was placed in-between the mice throughout each session allowing for sensory but never any physical contact. Each session lasted for 15min and was conducted every day for 16 consecutive days. After the termination of each session, mice were returned to their home cages to rest and consolidate the experience.

### RNA Isolation

Animals were sacrificed by isoflurane inhalation followed by decapitation, brains were extracted instantaneously followed by freezing in methylbutane and storage at −80°C. Frozen brains were sectioned in a cryostat microtome (Microm HM 560 M, ThermoScientific) at 100μm. Sectioning temperature of the knife was −10°C and of the specimen −11°C. The mPFC, blA, and vHC regions were punched with a brain punch tissue needle (Leica Biosystems) and stored at −80°C until processing. Total RNA was extracted according to the manufacturer’s protocol using RNeasy Micro Kit (Qiagen) in combination with TRIzol (Thermo Fisher Scientific). RNA was stored at −80 °C and freshly diluted for application.

Clontech’s SMART-Seq v4 Ultra Low Input RNA Kit (112219) was used for cDNA generation from 0,5ng of total RNA, according to the manufacturer’s protocol. cDNA was amplified by 16 cycles of LD-PCR. NGS library preparation was performed from 1ng of cDNA with Illumina`s Nextera XT DNA library prep Kit Reference Guide (May 2019, Document: 15031942v05), and amplified in 12 PCR cycles. Libraries were profiled in a High Sensitivity DNA Chip on a 2100 Bioanalyzer (Agilent Technologies) and quantified using the Qubit dsDNA HS Assay Kit, in a Qubit 2.0 Fluorometer (Life Technologies). The 81 samples (later referred to as trancriptome profiles) were pooled in equimolar ratio and sequenced on 8 NextSeq 500 High output Flowcells, SR for 1×75 cycles plus 2×8 cycles for the dual index read.

### Transcriptome Analysis

Raw sequencing read files of 81 transcriptome profiles were analysed with FastQC (version 0.11.8, www.bioinformatics.babraham.ac.uk/projects/fastqc/) and found to be of high quality (data not shown). Reads were mapped to the mouse genome (version GR-Cm38.p6) using the Star alignment software (version 2.5.3a, PMID 23104886) resulting in unique mapping rates >90% with ~ 35 mio mapped reads per sample on average. Transcript quantification was performed using the FeatureCounts software of the Subread package (version 1.5.3, PMID 24227677) and GENCODE mouse annotation (version M17, PMID 30357393). Linear and nonlinear dimension reduction for 2-d transcriptome profile representations were performed through PCA and t-SNE implementations in the R analysis environment. Outliers were identified by large distance in sample correlation and clustering plots as well as based on gene expression heatmaps resulting in the removal of 15 transcriptome profiles. Differential gene expression analysis between all groups was conducted using the DESeq2 R package (version 1.20.0, PMID 25516281). Only transcripts with at least 5 mapping reads in one of the corresponding transcriptome profiles were considered. DESeq2 with default parameters and the integrated log2-fold change shrinkage method ‘normal’ was used. Significantly differentially expressed genes were identified at adjusted p-value cutoff p-adjusted≤0.05. Intersecting and exclusive up-or down-regulated candidates between comparisons were identified and plotted as Venn diagrams using the R package “VennDiagram” (version 1.6.20). Metabolic pathway, gene ontology term and transcription factor enrichment were calculated using the Enrichr online tool (amp.phar-m.mssm.edu/Enrichr/, accessed March 2020, PMID 27141961). Volcano plots were produced using the EnhancedVolcano R package (version 1.4.0).

### Tracking

Video Tracking and automatic scoring of all behavioural tests were done using Ethovision software 11.0 by Noldus® (Wageningen, The Netherlands). All tests were corrected for nose-tail switches by a blind observer. In the case of Cue Discrimination Test and Active Avoidance Task, activity of mice was detected by infrared light beams.

### Behavioural Statistical Analysis

All statistical analysis was performed using GraphPad Prism software (version 6). Appropriate statistical tests were chosen based on the experimental conditions followed by outlier analysis using Grubb’s test with a significance level of 0.05 to exclude any run-away values. Similar variance between the groups being statistically compared was confirmed before conducting any analysis and all tests performed were two sided. When time was scored, it was calculated in percent of total time of the respective test (time%). Parametric statistical tests were always used. Exact sample number (n value), p-value, degrees of freedom, and F-value (in the case of ANOVAs) are all provided in the legend of the respective figure.

## DATA AVAILABILITY

The behavioural data that support the findings of this study are available from the corresponding author upon reasonable request. Raw RNA sequencing reads and gene counts are available on NCBI gene expression omnibus (GEO) under the accession number GSEXXXXX (provided before publication).

