## Supplementary Information for "Reconceptualising resilience within a translational framework is supported by unique and brain-region specific transcriptional signatures in mice"

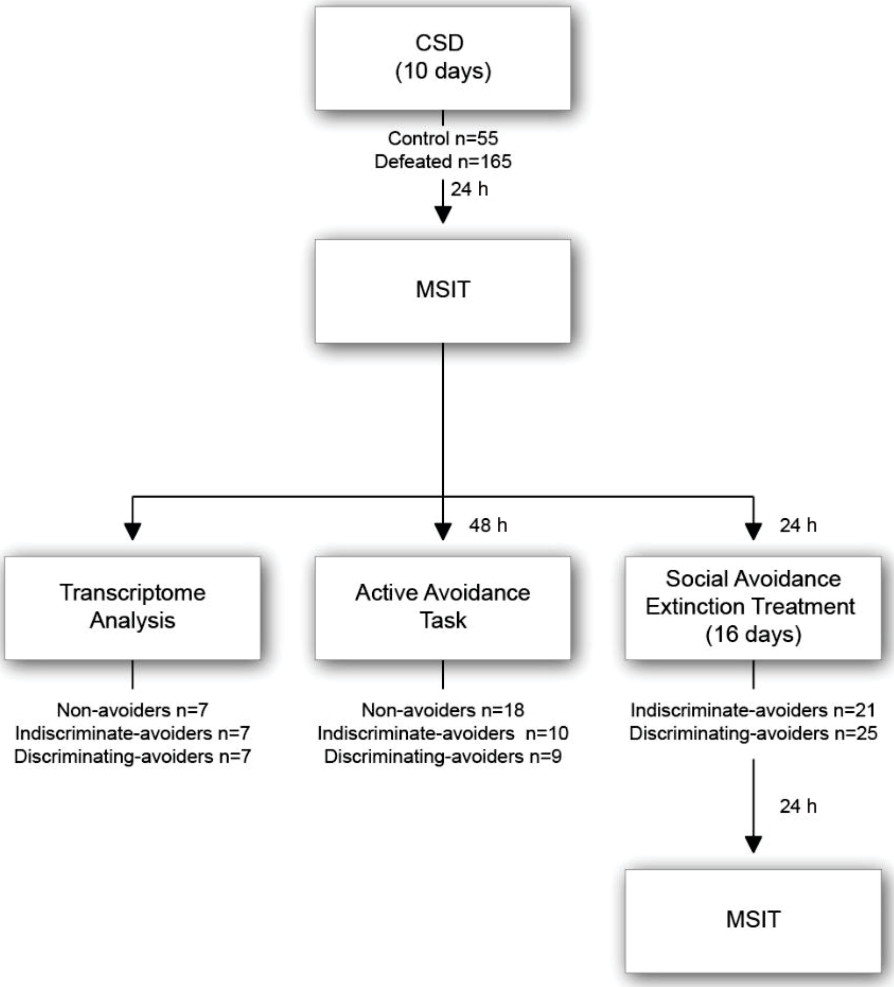
CSD
(10 days)
Control n=55
Defeated n=165
24 h
MSIT
48 h
24 h
Transcriptome Analysis
Active Avoidance Task
Social Avoidance Extinction Treatment (16 days)
Non-avoiders n=7
Indiscriminate-avoiders n=7
Discriminating-avoiders n=7
Non-avoiders n=18
Indiscriminate-avoiders n=10
Discriminating-avoiders n=9
Indiscriminate-avoiders n=21
Discriminating-avoiders n=25
24 h
MSIT

Figure S1. Schematic Timeline. Chronic social defeat (CSD) took place for 10 consecutive days on 193 mice (*Defeated*) while 43 animals served as a non-defeated control group. Following the last session by 24h, the Modified Social Interaction test (MSIT) was performed. Following the test, Defeated mice were divided into three subgroups in addition to the Control group. Transcriptome analysis was performed on 7 animals from each subgroup (in addition to the *Control* group) sacrificed immediately after the MSIT. Active avoidance task followed the MSIT 48h later on another subset of animals; n=18 *Non-avoiders*, n=10 *Indiscriminate-avoiders*, and n=9 *Discriminating-avoiders*. Social avoidance extinction treatment followed the MSIT 24h later and lasted for 16 consecutive days on yet another subset of animals; n=21 *Indiscriminate-avoiders* and n=25 *Discriminating-avoiders*. Following the last session by 24h, the second MSIT took place.

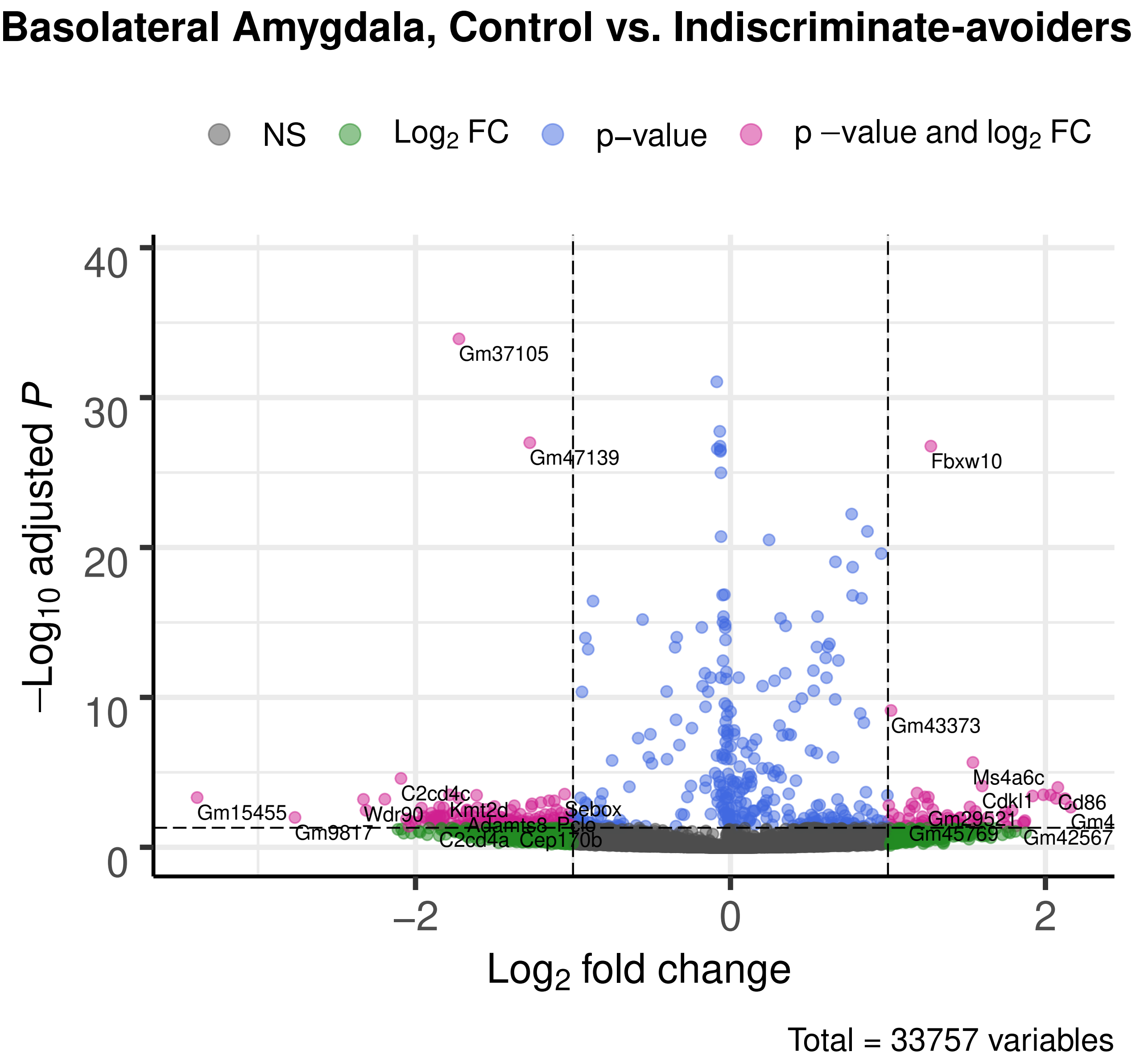
Basolateral Amygdala, Control vs. Indiscriminate-avoiders
NS
Log2 FC
p−value
p −value and log2 FC
−Log10 adjusted P
Log2 fold change
40
30
20
10
0
−2
0
2
Gm37105
Gm47139
Fbxw10
Gm43373
Ms4a6c
Cdkl1
Cd86
Gm4
Gm42567
Gm45769
Gm29521
Gm15455
Gm9817
Wdr90
C2cd4c
Kmt2d
Adamts8
Pclo
Sebox
C2cd4a
Cep170b
Total = 33757 variables

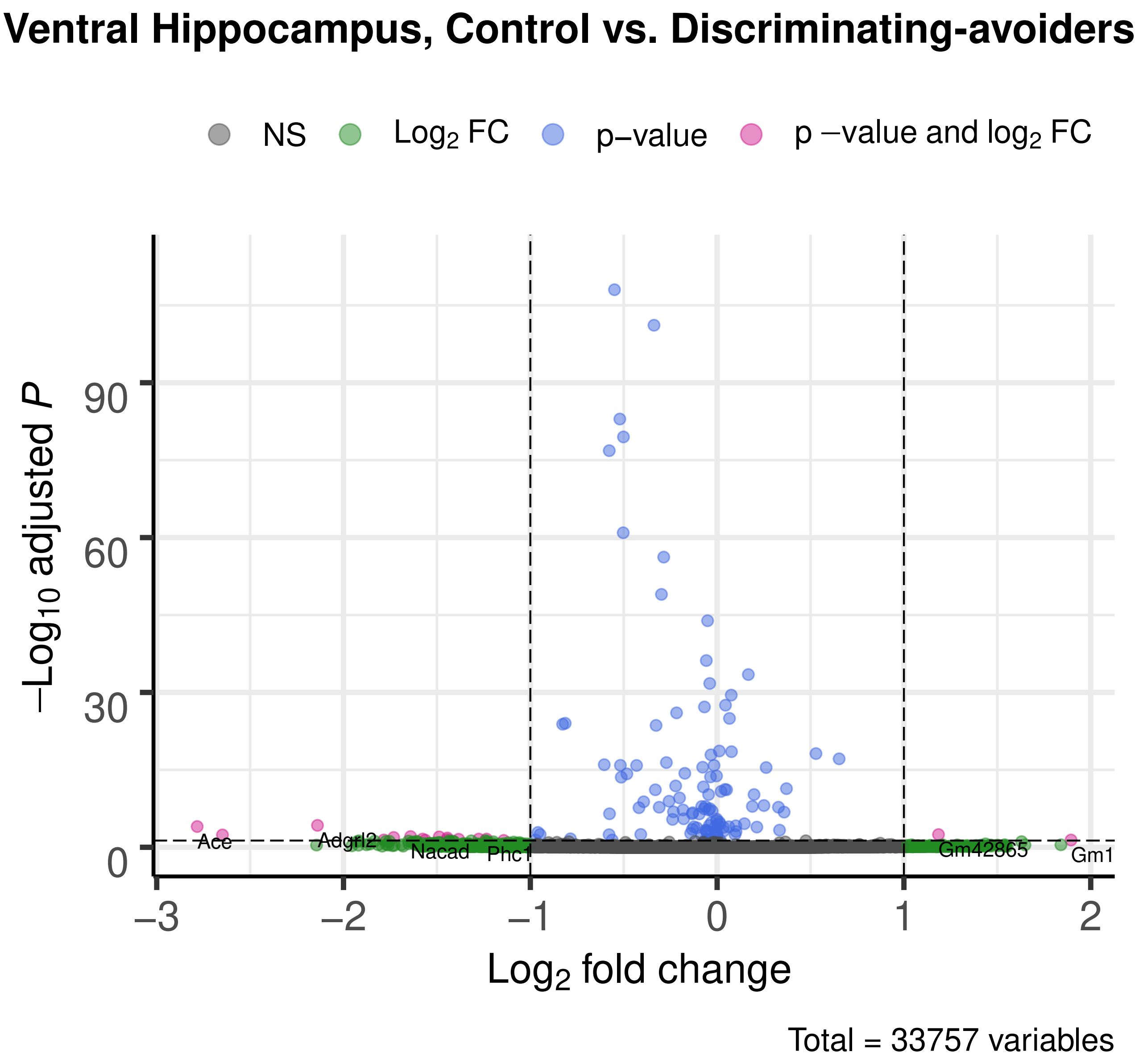

Ventral Hippocampus, Control vs. Discriminating-avoiders
NS
Log2 FC
p−value
p −value and log2 FC
−Log10 adjusted P
90
60
30
0
Ace
Adgrl2
Nacad
Phc1
Gm42865
Gm1
−3
−2
−1
0
1
2
Log2 fold change
Total = 33757 variables

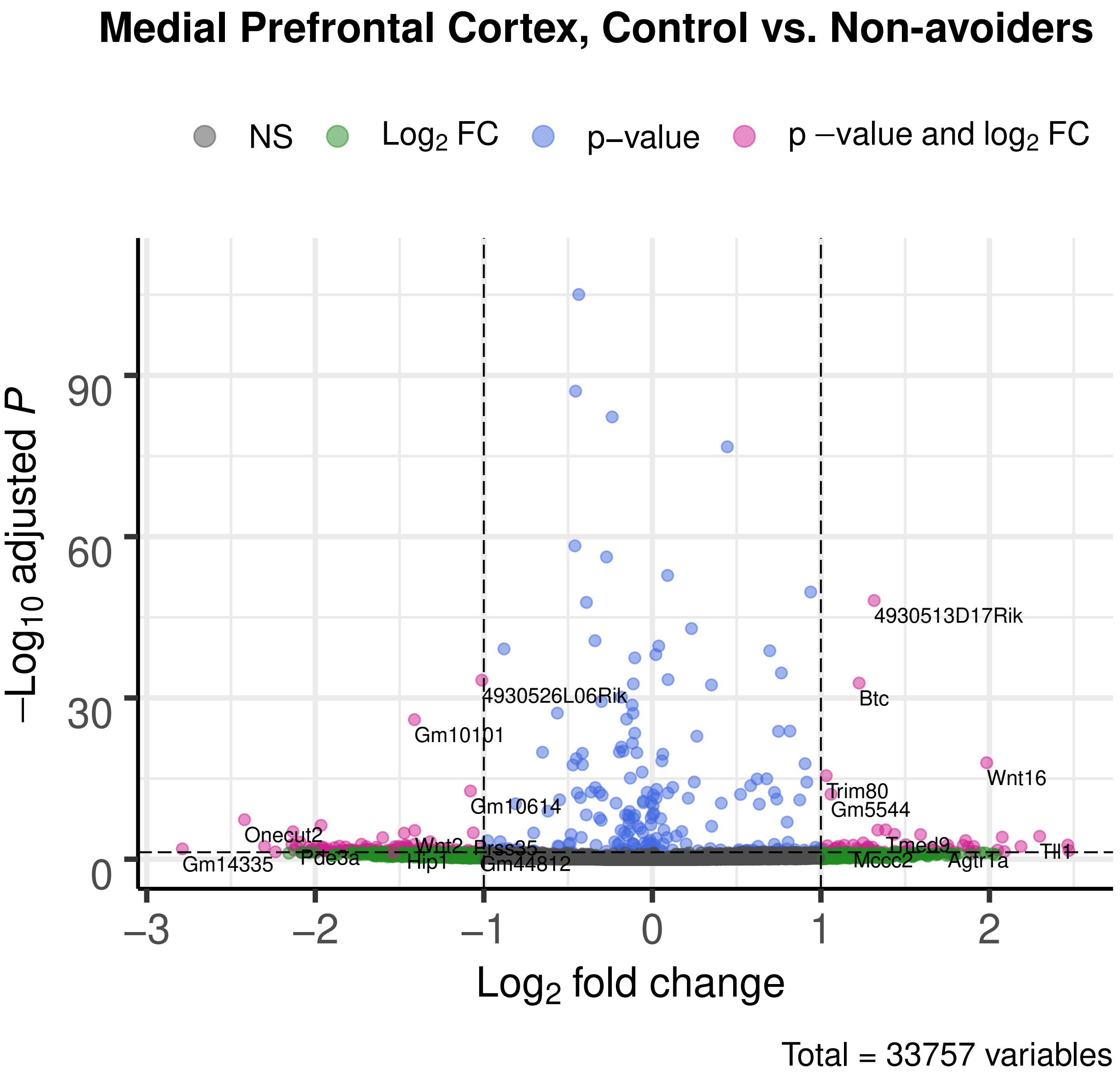
Medial Prefrontal Cortex, Control vs. Non-avoiders
NS
Log2 FC
p−value
p −value and log2 FC
−Log10 adjusted P
Log2 fold change
Total = 33757 variables
4930513D17Rik
Btc
4930526L06Rik
Gm10101
Wnt16
Trim80
Gm5544
Gm10614
Onecut2
Gm14335
Pde3a
Hip1
Wnt2
Prss35
Gm44812
Tmed9
Mccc2
Agtr1a
Tll1

Figure S2. Volcano Plot Analysis. Gene differential expression characteristics for: (a) The Basolateral Amygdala of the Indiscriminate-avoiders subgroup compared to non-defeated controls, (b) The Ventral Hippocampus of the Discriminating-avoiders subgroup compared to non-defeated controls, (c) The Medial Prefrontal Cortex of the Non-avoiders subgroup compared to non-defeated controls. The horizontal dashed line represents the adjusted p-value threshold (0.05) used for identifying differentially expresses genes. Analogously, the vertical dashed lines represent log2-fold-change threshold values (-1 and 1). Significantly (padj<0.05) and strongly (|log2fc|>1) differentially expressed genes are highlighted in magenta.

Table S1. Microarray Gene List. Differentially expressed genes (log2fc) between each of the three subgroups in comparison to the Control group in the three analysed brain regions; Basolateral Amygdala, Ventral Hippocampus, and Medial Prefrontal Cortex. Negative values indicate down-regulation whereas positive values indicate up-regulation.
