## Supplementary material for "Reconceptualising resilience within a translational framework is supported by unique and brain-region specific transcriptional signatures in mice": Table S1 Basolateral Amygdala

| | Discriminating-avoiders | | Indiscriminate-avoiders | | Non-avoiders | |
|---|---|---|---|---|---|---|
| **MGI symbol (Ensembl gene ID)** | *log2-fold-change* | *adjusted p-value* | *log2-fold-change* | *adjusted p-value* | *log2-fold-change* | *adjusted p-value* |
| 1500002F19Rik (ENSMUSG00000097595) | | | 1.18 | 0.0002 | | |
| 1700015F17Rik (ENSMUSG00000079666) | | | 1.58 | 0.0450 | | |
| 1700034G24Rik (ENSMUSG00000097962) | | | -1.42 | 0.0408 | | |
| 1700048F04Rik (ENSMUSG00000105274) | | | -1.02 | 0.0117 | | |
| 2500002B13Rik (ENSMUSG00000096917) | | | -1.68 | 0.0364 | | |
| 2900026A02Rik (ENSMUSG00000051339) | | | -1.76 | 0.0052 | | |
| 4732440D04Rik (ENSMUSG00000090031) | | | 1.38 | 0.0075 | | |
| 4930599N23Rik (ENSMUSG00000073144) | | | -1.21 | 0.0428 | | |
| 5031434O11Rik (ENSMUSG00000097885) | | | -1.29 | 0.0197 | | |
| A730071L15Rik (ENSMUSG00000096923) | | | -1.08 | 0.0266 | | |
| Aatk (ENSMUSG00000025375) | | | -1.44 | 0.0230 | | |
| Acap3 (ENSMUSG00000029033) | | | -1.38 | 0.0213 | | |
| Actg2 (ENSMUSG00000059430) | | | -1.01 | 0.0408 | | |
| Adamts2 (ENSMUSG00000036545) | | | -1.63 | 0.0399 | | |
| Adamts8 (ENSMUSG00000031994) | | | -1.67 | 0.0032 | | |
| Adgrb1 (ENSMUSG00000034730) | | | -1.37 | 0.0017 | | |
| Adgrb2 (ENSMUSG00000028782) | | | -1.35 | 0.0364 | | |
| Ak4 (ENSMUSG00000028527) | | | -1.26 | 0.0374 | | |
| Akr1cl (ENSMUSG00000025955) | | | 1.46 | 0.0362 | | |
| Alas2 (ENSMUSG00000025270) | | | 1.53 | 0.0354 | | |
| Alpk1 (ENSMUSG00000028028) | 1.26 | 0.0473 | | | | |
| Amigo1 (ENSMUSG00000050947) | | | -1.61 | 0.0023 | | |
| Ankmy1 (ENSMUSG00000034212) | | | -1.40 | 0.0139 | | |
| Anxa11 (ENSMUSG00000021866) | | | -1.76 | 0.0050 | | |
| Anxa11os (ENSMUSG00000098609) | | | -1.40 | 0.0162 | | |
| Ap3d1 (ENSMUSG00000020198) | | | -1.21 | 0.0266 | | |
| Arf4os (ENSMUSG00000090861) | | | 1.17 | 0.0020 | | |
| Arhgap23 (ENSMUSG00000049807) | | | -1.36 | 0.0207 | | |
| Arhgap35 (ENSMUSG00000058230) | | | -1.40 | 0.0034 | | |
| Asxl1 (ENSMUSG00000042548) | | | -1.22 | 0.0362 | | |
| Atrn (ENSMUSG00000027312) | | | -1.22 | 0.0189 | | |
| Avp (ENSMUSG00000037727) | -1.61 | 0.0473 | | | | |
| BE692007 (ENSMUSG00000099757) | 1.46 | 0.0002 | 1.23 | 0.0004 | 1.34 | 0.0007 |
| Birc3 (ENSMUSG00000032000) | 1.88 | 0.0034 | 2.07 | 0.0005 | | |
| Bmp3 (ENSMUSG00000029335) | | | | | | |
| Brsk2 (ENSMUSG00000053046) | | | -1.39 | 0.0318 | | |
| Bst1 (ENSMUSG00000029082) | | | 1.03 | 0.0454 | | |
| C030005K15Rik (ENSMUSG00000079183) | -1.01 | 0.0000 | | | | |
| C2cd4a (ENSMUSG00000047990) | | | -1.85 | 0.0307 | | |
| C2cd4c (ENSMUSG00000045912) | | | -2.09 | 0.0000 | | |
| Capg (ENSMUSG00000056737) | | | 1.73 | 0.0092 | | |
| Cavin2 (ENSMUSG00000045954) | | | 1.56 | 0.0040 | | |
| Cct6b (ENSMUSG00000020698) | | | | | 1.18 | 0.0409 |
| Cd86 (ENSMUSG00000022901) | 1.61 | 0.0159 | 2.08 | 0.0001 | 1.55 | 0.0413 |
| Cdc42bpb (ENSMUSG00000021279) | | | -1.56 | 0.0222 | | |
| Cdh24 (ENSMUSG00000059674) | | | -1.14 | 0.0062 | | |
| Cdk5r1 (ENSMUSG00000048895) | | | -1.20 | 0.0192 | | |
| Cdkl1 (ENSMUSG00000020990) | 1.23 | 0.0178 | 1.60 | 0.0001 | | |
| Cep170b (ENSMUSG00000072825) | | | -1.34 | 0.0308 | | |
| Ch25h (ENSMUSG00000050370) | | | 1.55 | 0.0091 | | |
| Chd5 (ENSMUSG00000005045) | | | -1.35 | 0.0463 | | |
| Cln5 (ENSMUSG00000022125) | | | 1.18 | 0.0364 | | |
| Clstn1 (ENSMUSG00000039953) | | | -1.54 | 0.0493 | | |
| Cmtm7 (ENSMUSG00000032436) | | | 1.32 | 0.0105 | | |
| Col22a1 (ENSMUSG00000079022) | | | -1.15 | 0.0008 | | |
| Cpeb4 (ENSMUSG00000020300) | | | -1.12 | 0.0008 | | |
| Crtc1 (ENSMUSG00000003575) | | | -1.51 | 0.0057 | | |
| Ctsc (ENSMUSG00000030560) | | | 1.29 | 0.0169 | | |
| D830036C21Rik (ENSMUSG00000108854) | | | 1.57 | 0.0489 | | |
| Dab2ip (ENSMUSG00000026883) | | | -1.12 | 0.0160 | | |
| Dcn (ENSMUSG00000019929) | | | | | 1.82 | 0.0174 |
| Defb42 (ENSMUSG00000054763) | | | 1.45 | 0.0255 | | |
| Dgkz (ENSMUSG00000040479) | | | -1.46 | 0.0224 | | |
| Dnajb5 (ENSMUSG00000036052) | | | -1.31 | 0.0424 | | |
| Dnajc6 (ENSMUSG00000028528) | | | -1.12 | 0.0381 | | |
| Dop1b (ENSMUSG00000022946) | | | -1.24 | 0.0339 | | |
| Dpysl2 (ENSMUSG00000022048) | | | -1.08 | 0.0263 | | |
| Dpysl4 (ENSMUSG00000025478) | | | -1.52 | 0.0092 | | |
| Emid1 (ENSMUSG00000034164) | | | -1.37 | 0.0204 | | |
| ENSMUSG00000097136 (ENSMUSG00000097136) | | | -2.04 | 0.0395 | | |
| ENSMUSG00000097272 (ENSMUSG00000097272) | | | -1.60 | 0.0214 | | |
| ENSMUSG00000097843 (ENSMUSG00000097843) | | | -1.82 | 0.0078 | | |
| ENSMUSG00000109782 (ENSMUSG00000109782) | | | 1.55 | 0.0195 | | |
| Epas1 (ENSMUSG00000024140) | | | -1.14 | 0.0489 | | |
| Epn2 (ENSMUSG00000001036) | | | -1.03 | 0.0339 | | |
| Esm1 (ENSMUSG00000042379) | 1.40 | 0.0444 | 1.21 | 0.0362 | | |
| Ets1 (ENSMUSG00000032035) | | | -1.57 | 0.0280 | | |
| Evc2 (ENSMUSG00000050248) | | | -1.95 | 0.0086 | | |
| Extl3 (ENSMUSG00000021978) | | | -1.22 | 0.0474 | | |
| Far2os2 (ENSMUSG00000086777) | | | -1.80 | 0.0479 | | |
| Fasn (ENSMUSG00000025153) | | | -1.30 | 0.0385 | | |
| Fbxl19 (ENSMUSG00000030811) | | | -1.56 | 0.0446 | | |
| Fbxw10 (ENSMUSG00000090173) | | | 1.27 | 0.0000 | | |
| Fermt1 (ENSMUSG00000027356) | | | -1.61 | 0.0003 | | |
| Fryl (ENSMUSG00000070733) | | | -1.07 | 0.0224 | | |
| Fsbp (ENSMUSG00000094595) | 1.34 | 0.0173 | 1.92 | 0.0004 | 1.22 | 0.0470 |
| Gli1 (ENSMUSG00000025407) | | | 1.74 | 0.0258 | | |
| Gm11192 (ENSMUSG00000086586) | | | -1.78 | 0.0274 | | |
| Gm11513 (ENSMUSG00000087630) | | | -1.03 | 0.0335 | | |
| Gm11733 (ENSMUSG00000069588) | 1.48 | 0.0121 | 1.23 | 0.0167 | 1.84 | 0.0026 |
| Gm12744 (ENSMUSG00000085023) | | | -1.73 | 0.0356 | | |
| Gm13607 (ENSMUSG00000081257) | 1.45 | 0.0207 | | | | |
| Gm13748 (ENSMUSG00000086836) | | | -1.07 | 0.0272 | | |
| Gm14327 (ENSMUSG00000074521) | | | 1.24 | 0.0383 | | |
| Gm14393 (ENSMUSG00000078905) | | | 1.87 | 0.0224 | | |
| Gm14439 (ENSMUSG00000084050) | 1.84 | 0.0041 | 2.12 | 0.0006 | 1.36 | 0.0500 |
| Gm15445 (ENSMUSG00000085311) | | | -2.31 | 0.0034 | | |
| Gm15455 (ENSMUSG00000081402) | -2.58 | 0.0321 | -3.39 | 0.0005 | -2.83 | 0.0160 |
| Gm15587 (ENSMUSG00000086769) | | | -1.65 | 0.0266 | | |
| Gm16316 (ENSMUSG00000087129) | | | -1.56 | 0.0186 | | |
| Gm16675 (ENSMUSG00000097534) | | | | | 1.11 | 0.0000 |
| Gm17039 (ENSMUSG00000092103) | | | -1.42 | 0.0392 | | |
| Gm18755 (ENSMUSG00000112712) | | | 1.40 | 0.0219 | | |
| Gm19557 (ENSMUSG00000097990) | 1.91 | 0.0019 | 1.74 | 0.0047 | 1.85 | 0.0054 |
| Gm20470 (ENSMUSG00000092190) | | | 1.18 | 0.0258 | | |
| Gm20655 (ENSMUSG00000093672) | 1.20 | 0.0017 | 1.15 | 0.0013 | 1.25 | 0.0021 |
| Gm21814 (ENSMUSG00000096299) | | | -1.28 | 0.0472 | | |
| Gm26587 (ENSMUSG00000097470) | | | -2.00 | 0.0224 | | |
| Gm26645 (ENSMUSG00000097684) | | | | | 1.12 | 0.0436 |
| Gm26708 (ENSMUSG00000097016) | | | 1.46 | 0.0111 | | |
| Gm26918 (ENSMUSG00000097976) | | | -1.12 | 0.0412 | | |
| Gm26953 (ENSMUSG00000097986) | | | -1.33 | 0.0062 | | |
| Gm27004 (ENSMUSG00000098027) | | | -1.15 | 0.0454 | | |
| Gm27761 (ENSMUSG00000098566) | -1.93 | 0.0347 | | | -2.24 | 0.0126 |
| Gm29460 (ENSMUSG00000099686) | | | 1.22 | 0.0213 | | |
| Gm29521 (ENSMUSG00000100807) | | | 1.25 | 0.0013 | | |
| Gm32816 (ENSMUSG00000108668) | | | | | -2.17 | 0.0500 |
| Gm32872 (ENSMUSG00000112793) | 1.77 | 0.0200 | | | | |
| Gm3375 (ENSMUSG00000107470) | | | 1.07 | 0.0499 | | |
| Gm37056 (ENSMUSG00000102753) | | | -1.25 | 0.0366 | | |
| Gm37105 (ENSMUSG00000102672) | | | -1.72 | 0.0000 | | |
| Gm37320 (ENSMUSG00000104324) | 1.30 | 0.0053 | | | 1.15 | 0.0166 |
| Gm37420 (ENSMUSG00000103869) | | | -1.59 | 0.0344 | | |
| Gm37499 (ENSMUSG00000103901) | | | -1.65 | 0.0296 | | |
| Gm37940 (ENSMUSG00000103899) | | | -1.96 | 0.0178 | | |
| Gm38043 (ENSMUSG00000104293) | | | 1.40 | 0.0215 | | |
| Gm38077 (ENSMUSG00000104344) | | | -2.19 | 0.0006 | | |
| Gm42567 (ENSMUSG00000105085) | | | 1.86 | 0.0218 | | |
| Gm42735 (ENSMUSG00000107041) | | | | | -1.90 | 0.0406 |
| Gm42875 (ENSMUSG00000105271) | | | -2.03 | 0.0190 | | |
| Gm43185 (ENSMUSG00000104621) | 1.12 | 0.0069 | 1.14 | 0.0036 | 1.21 | 0.0061 |
| Gm43331 (ENSMUSG00000104910) | | | -1.85 | 0.0021 | | |
| Gm43373 (ENSMUSG00000107363) | | | 1.02 | 0.0000 | | |
| Gm43437 (ENSMUSG00000104965) | | | 2.16 | 0.0021 | | |
| Gm43573 (ENSMUSG00000104786) | 1.91 | 0.0301 | 1.87 | 0.0167 | | |
| Gm44574 (ENSMUSG00000105311) | | | -1.87 | 0.0082 | | |
| Gm44698 (ENSMUSG00000109366) | | | 1.29 | 0.0497 | | |
| Gm45537 (ENSMUSG00000109606) | 1.44 | 0.0024 | | | | |
| Gm45769 (ENSMUSG00000107838) | 1.96 | 0.0003 | 1.13 | 0.0125 | 1.58 | 0.0036 |
| Gm47139 (ENSMUSG00000113106) | | | -1.27 | 0.0000 | | |
| Gm47246 (ENSMUSG00000114535) | | | -1.58 | 0.0305 | | |
| Gm49188 (ENSMUSG00000115300) | 1.29 | 0.0260 | 1.02 | 0.0325 | | |
| Gm5087 (ENSMUSG00000051729) | | | -1.84 | 0.0092 | | |
| Gm6352 (ENSMUSG00000107952) | 1.61 | 0.0401 | | | | |
| Gm8213 (ENSMUSG00000082474) | | | 1.76 | 0.0437 | | |
| Gm9169 (ENSMUSG00000098198) | | | 1.35 | 0.0393 | | |
| Gm9817 (ENSMUSG00000047061) | | | -2.77 | 0.0102 | | |
| Gm9877 (ENSMUSG00000052426) | | | 1.38 | 0.0111 | | |
| Gpc1 (ENSMUSG00000034220) | | | -1.55 | 0.0112 | | |
| Grik5 (ENSMUSG00000003378) | | | -1.09 | 0.0428 | | |
| Grk3 (ENSMUSG00000042249) | | | -1.01 | 0.0150 | | |
| Gse1 (ENSMUSG00000031822) | | | -1.42 | 0.0457 | | |
| Hba-a1 (ENSMUSG00000069919) | | | 1.57 | 0.0154 | | |
| Hba-a2 (ENSMUSG00000069917) | | | 1.46 | 0.0325 | | |
| Hbb-bs (ENSMUSG00000052305) | | | 1.63 | 0.0106 | | |
| Hbb-bt (ENSMUSG00000073940) | | | 1.79 | 0.0037 | | |
| Hectd4 (ENSMUSG00000042744) | | | -1.19 | 0.0415 | | |
| Hist1h1b (ENSMUSG00000058773) | | | 1.41 | 0.0149 | | |
| Hsd17b7 (ENSMUSG00000026675) | 1.10 | 0.0019 | | | | |
| I730030J21Rik (ENSMUSG00000116504) | | | 1.28 | 0.0031 | 1.58 | 0.0017 |
| Ift140 (ENSMUSG00000024169) | | | -1.46 | 0.0443 | | |
| Ighd (ENSMUSG00000104213) | 1.31 | 0.0021 | | | | |
| Igsf9b (ENSMUSG00000034275) | | | -1.45 | 0.0105 | | |
| Ildr2 (ENSMUSG00000040612) | | | -1.06 | 0.0477 | | |
| Ilrun (ENSMUSG00000056692) | | | -1.02 | 0.0329 | | |
| Ism1 (ENSMUSG00000074766) | | | | | 1.51 | 0.0431 |
| Khsrp (ENSMUSG00000007670) | | | -1.57 | 0.0450 | | |
| Kif3b (ENSMUSG00000027475) | | | -1.18 | 0.0065 | | |
| Kmt2d (ENSMUSG00000048154) | | | -1.78 | 0.0003 | | |
| Kpnb1 (ENSMUSG00000001440) | | | -1.08 | 0.0109 | | |
| Leng8 (ENSMUSG00000035545) | | | -1.30 | 0.0213 | | |
| Lingo1 (ENSMUSG00000049556) | | | -1.69 | 0.0111 | | |
| Lrfn1 (ENSMUSG00000030600) | | | -1.94 | 0.0111 | | |
| Lrrc8a (ENSMUSG00000007476) | | | -1.31 | 0.0310 | | |
| Ltbp4 (ENSMUSG00000040488) | | | -1.82 | 0.0105 | | |
| Madd (ENSMUSG00000040687) | | | -1.20 | 0.0499 | | |
| Mast2 (ENSMUSG00000003810) | | | -1.25 | 0.0310 | | |
| Mef2d (ENSMUSG00000001419) | | | -1.33 | 0.0356 | | |
| Meox1 (ENSMUSG00000001493) | | | 1.42 | 0.0215 | | |
| Mettl18 (ENSMUSG00000041396) | | | 1.07 | 0.0111 | | |
| Mettl21c (ENSMUSG00000047343) | | | -2.06 | 0.0131 | | |
| Mfsd12 (ENSMUSG00000034854) | | | -1.34 | 0.0367 | | |
| Mical2 (ENSMUSG00000038244) | | | -1.71 | 0.0111 | | |
| Mir7055 (ENSMUSG00000106470) | | | -1.86 | 0.0084 | | |
| Mis18bp1 (ENSMUSG00000047534) | | | 1.01 | 0.0016 | | |
| Mmp23 (ENSMUSG00000029061) | | | | | 1.01 | 0.0321 |
| Mrtfa (ENSMUSG00000042292) | | | -1.59 | 0.0329 | | |
| Mrvi1 (ENSMUSG00000005611) | | | | | 1.01 | 0.0357 |
| Ms4a4a (ENSMUSG00000101389) | 1.12 | 0.0017 | 1.26 | 0.0005 | 1.03 | 0.0046 |
| Ms4a6c (ENSMUSG00000079419) | 1.11 | 0.0001 | 1.54 | 0.0000 | 1.15 | 0.0001 |
| Myh6 (ENSMUSG00000040752) | | | -1.89 | 0.0272 | | |
| Nav1 (ENSMUSG00000009418) | | | -1.14 | 0.0428 | | |
| Nepro (ENSMUSG00000036208) | | | -1.05 | 0.0373 | | |
| Neurod1 (ENSMUSG00000034701) | | | -1.47 | 0.0288 | | |
| Nkd1 (ENSMUSG00000031661) | | | -1.18 | 0.0489 | | |
| Nptx1 (ENSMUSG00000025582) | | | -1.67 | 0.0304 | | |
| Nptxr (ENSMUSG00000022421) | | | -1.88 | 0.0140 | | |
| Ntf5 (ENSMUSG00000074121) | | | -1.24 | 0.0129 | | |
| Ntng2 (ENSMUSG00000035513) | | | -1.82 | 0.0091 | | |
| Pabpc4l (ENSMUSG00000090919) | 1.40 | 0.0091 | 1.99 | 0.0003 | | |
| Pcdha1 (ENSMUSG00000103442) | | | -1.13 | 0.0494 | | |
| Pcdha12 (ENSMUSG00000103310) | | | -1.17 | 0.0424 | | |
| Pcdha2 (ENSMUSG00000104148) | | | -1.20 | 0.0345 | | |
| Pcdha3 (ENSMUSG00000102312) | | | -1.18 | 0.0356 | | |
| Pcdha4 (ENSMUSG00000104252) | | | -1.17 | 0.0409 | | |
| Pcdha6 (ENSMUSG00000103707) | | | -1.14 | 0.0476 | | |
| Pcdha7 (ENSMUSG00000104318) | | | -1.17 | 0.0381 | | |
| Pcdha8 (ENSMUSG00000103800) | | | -1.11 | 0.0468 | | |
| Pcdha9 (ENSMUSG00000103770) | | | -1.21 | 0.0310 | | |
| Pcdhac1 (ENSMUSG00000103255) | | | -1.18 | 0.0367 | | |
| Pclo (ENSMUSG00000061601) | | | -1.10 | 0.0031 | | |
| Pip4k2c (ENSMUSG00000025417) | | | -1.44 | 0.0139 | | |
| Pip5k1c (ENSMUSG00000034902) | | | -1.38 | 0.0489 | | |
| Plekha6 (ENSMUSG00000041757) | | | -1.55 | 0.0122 | | |
| Pnlip (ENSMUSG00000046008) | | | 1.45 | 0.0425 | | |
| Polg2 (ENSMUSG00000020718) | | | 1.52 | 0.0020 | | |
| Ppfia3 (ENSMUSG00000003863) | | | -1.72 | 0.0005 | | |
| Prkcg (ENSMUSG00000078816) | | | -1.74 | 0.0356 | | |
| Prrt3 (ENSMUSG00000045009) | | | -1.28 | 0.0400 | | |
| Rab11fip4 (ENSMUSG00000017639) | | | -1.01 | 0.0336 | | |
| Rai1 (ENSMUSG00000062115) | | | -1.55 | 0.0030 | | |
| Rimbp2 (ENSMUSG00000029420) | | | -1.61 | 0.0189 | | |
| Rimbp3 (ENSMUSG00000071636) | | | -1.07 | 0.0051 | | |
| Rnf157 (ENSMUSG00000052949) | | | -1.25 | 0.0231 | | |
| Rtn4r (ENSMUSG00000043811) | | | -1.90 | 0.0102 | | |
| Rtn4rl2 (ENSMUSG00000050896) | | | -1.69 | 0.0408 | | |
| Rusc2 (ENSMUSG00000035969) | | | -1.51 | 0.0232 | | |
| Satb1 (ENSMUSG00000023927) | | | -1.37 | 0.0420 | | |
| Sbf1 (ENSMUSG00000036529) | | | -1.67 | 0.0037 | | |
| Sebox (ENSMUSG00000001103) | | | -1.05 | 0.0003 | | |
| Sema7a (ENSMUSG00000038264) | | | -1.82 | 0.0364 | | |
| Setd7 (ENSMUSG00000037111) | | | -1.16 | 0.0145 | | |
| Shank2 (ENSMUSG00000037541) | | | -1.33 | 0.0169 | | |
| Skida1 (ENSMUSG00000054074) | | | -1.43 | 0.0381 | | |
| Slc15a3 (ENSMUSG00000024737) | | | -1.44 | 0.0286 | | |
| Slc17a7 (ENSMUSG00000070570) | | | -1.90 | 0.0053 | | |
| Slc23a1 (ENSMUSG00000024354) | | | 1.13 | 0.0416 | | |
| Slc25a22 (ENSMUSG00000019082) | | | -1.35 | 0.0458 | | |
| Slurp1 (ENSMUSG00000022596) | | | 1.81 | 0.0374 | | |
| Sobp (ENSMUSG00000038248) | | | -1.18 | 0.0468 | | |
| Spred3 (ENSMUSG00000037239) | | | -1.46 | 0.0095 | | |
| Srcin1 (ENSMUSG00000038453) | | | -1.19 | 0.0195 | | |
| Srgap2 (ENSMUSG00000026425) | | | -1.02 | 0.0393 | | |
| Stn1 (ENSMUSG00000042694) | 1.10 | 0.0025 | | | | |
| Synpo (ENSMUSG00000043079) | | | -1.66 | 0.0107 | | |
| Synpr (ENSMUSG00000056296) | 1.21 | 0.0368 | 1.11 | 0.0364 | | |
| Tbc1d24 (ENSMUSG00000036473) | | | -1.50 | 0.0019 | | |
| Tbr1 (ENSMUSG00000035033) | | | -1.90 | 0.0195 | | |
| Tcof1 (ENSMUSG00000024613) | | | -1.96 | 0.0024 | | |
| Tead3 (ENSMUSG00000002249) | | | | | 2.37 | 0.0023 |
| Tecpr1 (ENSMUSG00000066621) | | | -1.30 | 0.0248 | | |
| Tecpr2 (ENSMUSG00000021275) | | | -1.36 | 0.0499 | | |
| Tent4a (ENSMUSG00000034575) | | | -1.58 | 0.0102 | | |
| Thsd4 (ENSMUSG00000032289) | | | -1.86 | 0.0213 | | |
| Ticam2 (ENSMUSG00000056130) | | | 1.03 | 0.0063 | | |
| Tmem132a (ENSMUSG00000024736) | | | -1.64 | 0.0077 | | |
| Tmem8b (ENSMUSG00000078716) | | | -1.37 | 0.0100 | | |
| Tnpo2 (ENSMUSG00000031691) | | | -1.12 | 0.0111 | | |
| Tnr (ENSMUSG00000015829) | | | -1.20 | 0.0369 | | |
| Topbp1 (ENSMUSG00000032555) | | | -1.21 | 0.0096 | | |
| Trak1 (ENSMUSG00000032536) | | | -1.20 | 0.0291 | | |
| Trappc10 (ENSMUSG00000000374) | | | -1.26 | 0.0013 | | |
| Trdc (ENSMUSG00000104876) | | | 1.00 | 0.0168 | | |
| Trim30a (ENSMUSG00000030921) | | | 1.45 | 0.0399 | | |
| Trrap (ENSMUSG00000045482) | | | -1.08 | 0.0222 | | |
| Ube2l6 (ENSMUSG00000027078) | | | 1.32 | 0.0325 | | |
| Unc13d (ENSMUSG00000057948) | 1.48 | 0.0050 | 1.22 | 0.0091 | | |
| Upf1 (ENSMUSG00000058301) | | | -1.57 | 0.0469 | | |
| Usp24 (ENSMUSG00000028514) | | | -1.02 | 0.0058 | | |
| Usp5 (ENSMUSG00000038429) | | | -1.12 | 0.0364 | | |
| Vars (ENSMUSG00000007029) | | | -1.47 | 0.0413 | | |
| Vmn1r4 (ENSMUSG00000071893) | 1.02 | 0.0235 | | | | |
| Vps13c (ENSMUSG00000035284) | -1.25 | 0.0009 | -1.20 | 0.0010 | -1.31 | 0.0008 |
| Vwa5b2 (ENSMUSG00000046613) | | | -1.62 | 0.0112 | | |
| Wdr90 (ENSMUSG00000073434) | | | -2.33 | 0.0006 | | |
| Ywhag (ENSMUSG00000051391) | | | -1.34 | 0.0452 | | |
| Zbtb33 (ENSMUSG00000048047) | | | -1.85 | 0.0077 | | |
| Zbtb8a (ENSMUSG00000028807) | | | 1.46 | 0.0107 | | |
| Zdhhc8 (ENSMUSG00000060166) | | | -1.82 | 0.0023 | | |
| Zfp839 (ENSMUSG00000021271) | | | -1.54 | 0.0232 | | |
| Zfp977 (ENSMUSG00000092335) | 1.71 | 0.0052 | 2.03 | 0.0003 | 1.56 | 0.0255 |
| Zmat3 (ENSMUSG00000027663) | | | -1.26 | 0.0019 | | |
| Zmiz1 (ENSMUSG00000007817) | | | -1.29 | 0.0121 | | |
| Zswim8 (ENSMUSG00000021819) | | | -1.43 | 0.0241 | | |
