## Supplementary material for "Reconceptualising resilience within a translational framework is supported by unique and brain-region specific transcriptional signatures in mice": Table S1 Medial Prefrontal Cortex

| | Non-avoiders | | Indiscriminate-avoiders | | Discriminating-avoiders | |
|---|---|---|---|---|---|---|
| **MGI symbol (Ensembl gene ID)** | *log2-fold-change* | *adjusted p-value* | *log2-fold-change* | *adjusted p-value* | *log2-fold-change* | *adjusted p-value* |
| 1700041G16Rik (ENSMUSG00000054822) | 2.46 | 0.0023 | | | | |
| 1700123O21Rik (ENSMUSG00000101481) | -1.55 | 0.0476 | | | | |
| 4833418N02Rik (ENSMUSG00000085287) | 1.02 | 0.0146 | 1.34 | 0.0029 | | |
| 4930451G09Rik (ENSMUSG00000022543) | -1.97 | 0.0129 | | | | |
| 4930513D17Rik (ENSMUSG00000106741) | 1.32 | 0.0000 | | | | |
| 4930513L16Rik (ENSMUSG00000116491) | -1.68 | 0.0156 | | | | |
| 4930526L06Rik (ENSMUSG00000097091) | -1.01 | 0.0000 | -1.06 | 0.0000 | | |
| 4930593A02Rik (ENSMUSG00000097725) | 1.87 | 0.0054 | 1.40 | 0.0283 | | |
| 4930599N23Rik (ENSMUSG00000073144) | -1.40 | 0.0298 | | | | |
| 4933407K13Rik (ENSMUSG00000087396) | -1.64 | 0.0450 | | | | |
| 4933440N22Rik (ENSMUSG00000042087) | 1.07 | 0.0296 | | | | |
| 5730405O15Rik (ENSMUSG00000087424) | -1.66 | 0.0496 | | | | |
| 6720483E21Rik (ENSMUSG00000097934) | -1.82 | 0.0052 | | | | |
| 9330158H04Rik (ENSMUSG00000073154) | -1.85 | 0.0244 | | | | |
| 9330160F10Rik (ENSMUSG00000072809) | 1.86 | 0.0003 | | | | |
| 9530046B11Rik (ENSMUSG00000086047) | -1.78 | 0.0328 | | | | |
| A330023F24Rik (ENSMUSG00000096929) | -1.34 | 0.0386 | | | | |
| A330070K13Rik (ENSMUSG00000052014) | -1.28 | 0.0261 | | | | |
| A730020M07Rik (ENSMUSG00000044522) | | | 1.08 | 0.0000 | | |
| Acer2 (ENSMUSG00000038007) | | | 1.51 | 0.0449 | | |
| Acot10 (ENSMUSG00000047565) | -1.71 | 0.0388 | -1.80 | 0.0161 | | |
| Aen (ENSMUSG00000030609) | 1.34 | 0.0000 | | | | |
| Ago3 (ENSMUSG00000028842) | -1.20 | 0.0379 | | | | |
| Agtr1a (ENSMUSG00000049115) | 1.75 | 0.0023 | | | | |
| Airn (ENSMUSG00000078247) | -1.60 | 0.0001 | | | | |
| Alas2 (ENSMUSG00000025270) | | | 2.42 | 0.0004 | | |
| Amt (ENSMUSG00000032607) | -1.64 | 0.0061 | | | | |
| Ankrd1 (ENSMUSG00000024803) | | | | | 1.81 | 0.0000 |
| Anxa1 (ENSMUSG00000024659) | | | 2.70 | 0.0040 | | |
| Anxa4 (ENSMUSG00000029994) | | | 1.72 | 0.0297 | | |
| Apoc2 (ENSMUSG00000002992) | | | 1.21 | 0.0000 | | |
| Arhgap9 (ENSMUSG00000040345) | 1.61 | 0.0440 | | | | |
| Ascl2 (ENSMUSG00000009248) | | | -1.36 | 0.0319 | | |
| Asf1b (ENSMUSG00000005470) | | | 1.36 | 0.0281 | | |
| Atp13a3 (ENSMUSG00000022533) | -1.38 | 0.0253 | | | | |
| Atp7a (ENSMUSG00000033792) | 1.91 | 0.0379 | | | | |
| Atxn1 (ENSMUSG00000046876) | -1.52 | 0.0118 | | | | |
| Avpr2 (ENSMUSG00000031390) | -2.03 | 0.0307 | | | | |
| B230104I21Rik (ENSMUSG00000097871) | 1.69 | 0.0426 | | | | |
| B430219N15Rik (ENSMUSG00000085211) | | | 1.21 | 0.0348 | | |
| BC016548 (ENSMUSG00000070794) | 1.44 | 0.0326 | | | | |
| Brsk1 (ENSMUSG00000035390) | -1.63 | 0.0172 | | | | |
| Btc (ENSMUSG00000082361) | 1.23 | 0.0000 | | | | |
| C030017D09Rik (ENSMUSG00000090907) | -1.76 | 0.0326 | | | | |
| C130060C02Rik (ENSMUSG00000097557) | -1.78 | 0.0417 | | | | |
| C1ra (ENSMUSG00000055172) | | | 1.11 | 0.0215 | | |
| C2 (ENSMUSG00000024371) | | | 1.99 | 0.0055 | | |
| C2cd4d (ENSMUSG00000091648) | -1.81 | 0.0211 | | | | |
| C630043F03Rik (ENSMUSG00000084910) | | | 1.04 | 0.0132 | | |
| CAAA01118383.1 (ENSMUSG00000063897) | -1.11 | 0.0382 | | | | |
| Cav1 (ENSMUSG00000007655) | | | 1.41 | 0.0204 | | |
| Ccl12 (ENSMUSG00000035352) | | | 2.66 | 0.0067 | | |
| Cd27 (ENSMUSG00000030336) | -1.96 | 0.0234 | | | | |
| Cd300ld (ENSMUSG00000034641) | | | 1.18 | 0.0000 | | |
| Cdc20 (ENSMUSG00000006398) | | | 1.23 | 0.0004 | | |
| Cdr1 (ENSMUSG00000090546) | -1.55 | 0.0176 | | | | |
| Cfap57 (ENSMUSG00000028730) | 1.50 | 0.0386 | | | | |
| Cgn (ENSMUSG00000068876) | -1.79 | 0.0177 | | | | |
| Ciart (ENSMUSG00000038550) | 1.09 | 0.0095 | 1.04 | 0.0112 | | |
| Cmtm3 (ENSMUSG00000031875) | | | 1.19 | 0.0368 | | |
| Crlf3 (ENSMUSG00000017561) | | | 1.06 | 0.0256 | | |
| Ctsc (ENSMUSG00000030560) | | | 1.91 | 0.0001 | | |
| Cubn (ENSMUSG00000026726) | | | 1.83 | 0.0262 | | |
| Cyp4f18 (ENSMUSG00000003484) | | | 2.57 | 0.0103 | | |
| Cysltr2 (ENSMUSG00000033470) | | | | | 1.03 | 0.0000 |
| D230049E03Rik (ENSMUSG00000113196) | | | | | -1.64 | 0.0009 |
| D5Ertd579e (ENSMUSG00000029190) | -1.35 | 0.0190 | | | | |
| D7Ertd443e (ENSMUSG00000030994) | | | -1.89 | 0.0138 | | |
| D930007J09Rik (ENSMUSG00000042874) | 1.15 | 0.0370 | | | | |
| Dars2 (ENSMUSG00000026709) | | | | | 1.05 | 0.0425 |
| Dennd4a (ENSMUSG00000053641) | -1.21 | 0.0382 | | | | |
| Dgka (ENSMUSG00000025357) | -1.71 | 0.0316 | | | | |
| Dnph1 (ENSMUSG00000040658) | 1.33 | 0.0270 | | | | |
| E330020D12Rik (ENSMUSG00000073538) | -1.94 | 0.0224 | | | | |
| ENSMUSG00000096954 (ENSMUSG00000096954) | -1.54 | 0.0488 | | | | |
| ENSMUSG00000097423 (ENSMUSG00000097423) | | | | | -1.39 | 0.0121 |
| Etv4 (ENSMUSG00000017724) | -1.86 | 0.0349 | | | | |
| Evi2b (ENSMUSG00000093938) | 1.91 | 0.0042 | 1.92 | 0.0025 | 1.70 | 0.0261 |
| Fads6 (ENSMUSG00000044788) | -1.78 | 0.0244 | | | | |
| Fam193b (ENSMUSG00000021495) | -1.46 | 0.0023 | | | | |
| Fam221a (ENSMUSG00000047115) | | | 1.37 | 0.0000 | 1.33 | 0.0001 |
| Far2os2 (ENSMUSG00000086777) | -1.92 | 0.0303 | | | | |
| Fbp1 (ENSMUSG00000069805) | | | 1.09 | 0.0446 | | |
| Fen1 (ENSMUSG00000024742) | 1.31 | 0.0057 | | | | |
| Fgl1 (ENSMUSG00000031594) | 1.10 | 0.0041 | 1.69 | 0.0001 | 1.79 | 0.0002 |
| Fgl2 (ENSMUSG00000039899) | | | 1.89 | 0.0211 | | |
| Fibin (ENSMUSG00000074971) | | | 1.74 | 0.0484 | | |
| Fxyd5 (ENSMUSG00000009687) | | | 1.23 | 0.0397 | | |
| Gad1os (ENSMUSG00000087264) | | | 1.85 | 0.0018 | | |
| Gm10101 (ENSMUSG00000061510) | -1.41 | 0.0000 | | | -1.29 | 0.0000 |
| Gm10143 (ENSMUSG00000064032) | 1.28 | 0.0076 | | | | |
| Gm10614 (ENSMUSG00000097773) | -1.08 | 0.0000 | -1.14 | 0.0000 | | |
| Gm10652 (ENSMUSG00000106329) | -1.21 | 0.0076 | | | | |
| Gm11201 (ENSMUSG00000085941) | | | 1.47 | 0.0468 | | |
| Gm11831 (ENSMUSG00000086625) | | | | | -1.21 | 0.0229 |
| Gm12098 (ENSMUSG00000082078) | 2.05 | 0.0203 | | | | |
| Gm12940 (ENSMUSG00000085334) | -1.32 | 0.0006 | | | | |
| Gm13483 (ENSMUSG00000085862) | | | | | -1.96 | 0.0270 |
| Gm13556 (ENSMUSG00000087294) | -1.88 | 0.0069 | | | | |
| Gm13589 (ENSMUSG00000085950) | 1.21 | 0.0089 | 1.29 | 0.0045 | | |
| Gm14335 (ENSMUSG00000081805) | -2.79 | 0.0119 | -2.86 | 0.0051 | -2.88 | 0.0122 |
| Gm15328 (ENSMUSG00000086095) | -1.81 | 0.0222 | | | | |
| Gm15385 (ENSMUSG00000083093) | | | -2.07 | 0.0394 | | |
| Gm15413 (ENSMUSG00000053049) | -1.71 | 0.0258 | | | | |
| Gm15422 (ENSMUSG00000082738) | | | -1.62 | 0.0373 | | |
| Gm16151 (ENSMUSG00000085280) | -1.65 | 0.0208 | | | | |
| Gm17195 (ENSMUSG00000091534) | | | 1.16 | 0.0330 | | |
| Gm17530 (ENSMUSG00000103731) | -1.65 | 0.0301 | | | | |
| Gm17938 (ENSMUSG00000114603) | | | -2.35 | 0.0345 | | |
| Gm18194 (ENSMUSG00000109724) | | | 1.98 | 0.0215 | | |
| Gm18782 (ENSMUSG00000109367) | | | 1.57 | 0.0014 | 1.21 | 0.0136 |
| Gm18956 (ENSMUSG00000102851) | -1.73 | 0.0016 | | | | |
| Gm19031 (ENSMUSG00000093748) | | | 1.08 | 0.0484 | | |
| Gm19605 (ENSMUSG00000113585) | -1.83 | 0.0392 | | | | |
| Gm20045 (ENSMUSG00000103983) | -1.98 | 0.0219 | | | | |
| Gm22489 (ENSMUSG00000080463) | 1.65 | 0.0116 | 1.56 | 0.0147 | | |
| Gm22879 (ENSMUSG00000064829) | | | -1.95 | 0.0454 | | |
| Gm23134 (ENSMUSG00000077990) | | | 1.16 | 0.0478 | | |
| Gm25328 (ENSMUSG00000095087) | -1.89 | 0.0379 | | | | |
| Gm26575 (ENSMUSG00000097926) | -2.02 | 0.0134 | | | | |
| Gm26747 (ENSMUSG00000097783) | 1.12 | 0.0071 | | | | |
| Gm26839 (ENSMUSG00000097917) | | | 1.18 | 0.0392 | | |
| Gm26974 (ENSMUSG00000098160) | -1.46 | 0.0061 | | | | |
| Gm27151 (ENSMUSG00000098739) | -1.73 | 0.0275 | | | | |
| Gm27206 (ENSMUSG00000098889) | -1.61 | 0.0460 | | | | |
| Gm27252 (ENSMUSG00000098708) | -1.86 | 0.0037 | | | | |
| Gm28154 (ENSMUSG00000100053) | -2.04 | 0.0063 | | | | |
| Gm29083 (ENSMUSG00000100811) | -1.96 | 0.0051 | | | | |
| Gm29502 (ENSMUSG00000100596) | | | | | 1.64 | 0.0236 |
| Gm29650 (ENSMUSG00000099876) | -2.30 | 0.0042 | | | | |
| Gm31166 (ENSMUSG00000109695) | -1.41 | 0.0016 | | | | |
| Gm32442 (ENSMUSG00000112404) | | | | | 1.47 | 0.0257 |
| Gm3325 (ENSMUSG00000113836) | | | 1.42 | 0.0067 | | |
| Gm3365 (ENSMUSG00000111594) | -1.70 | 0.0132 | | | | |
| Gm33680 (ENSMUSG00000112054) | | | 1.11 | 0.0042 | 1.44 | 0.0021 |
| Gm34934 (ENSMUSG00000115100) | -1.07 | 0.0496 | | | | |
| Gm35501 (ENSMUSG00000111068) | | | | | -1.69 | 0.0159 |
| Gm36236 (ENSMUSG00000113774) | -1.39 | 0.0297 | | | | |
| Gm36930 (ENSMUSG00000103778) | -2.08 | 0.0484 | | | | |
| Gm37069 (ENSMUSG00000104415) | -1.24 | 0.0100 | | | | |
| Gm37720 (ENSMUSG00000103959) | -2.00 | 0.0399 | | | | |
| Gm37814 (ENSMUSG00000102816) | | | -1.28 | 0.0297 | | |
| Gm38020 (ENSMUSG00000103697) | -1.94 | 0.0211 | | | | |
| Gm39323 (ENSMUSG00000111229) | | | | | -1.64 | 0.0329 |
| Gm4221 (ENSMUSG00000096948) | -1.65 | 0.0437 | | | | |
| Gm42495 (ENSMUSG00000106275) | 1.50 | 0.0454 | | | | |
| Gm42500 (ENSMUSG00000104727) | 1.50 | 0.0020 | | | | |
| Gm42588 (ENSMUSG00000105584) | | | -1.15 | 0.0000 | | |
| Gm42728 (ENSMUSG00000107158) | | | | | -1.31 | 0.0000 |
| Gm42732 (ENSMUSG00000107331) | | | | | -1.31 | 0.0288 |
| Gm42875 (ENSMUSG00000105271) | -2.03 | 0.0389 | | | | |
| Gm42928 (ENSMUSG00000105813) | | | 1.01 | 0.0130 | | |
| Gm43032 (ENSMUSG00000106498) | -1.05 | 0.0416 | | | | |
| Gm43200 (ENSMUSG00000105457) | -1.57 | 0.0224 | | | | |
| Gm43338 (ENSMUSG00000105679) | -2.00 | 0.0089 | | | | |
| Gm43376 (ENSMUSG00000104677) | -1.99 | 0.0262 | | | | |
| Gm43413 (ENSMUSG00000104951) | -1.78 | 0.0273 | | | | |
| Gm43519 (ENSMUSG00000105528) | -1.70 | 0.0137 | | | | |
| Gm4353 (ENSMUSG00000091900) | 2.09 | 0.0375 | | | | |
| Gm43628 (ENSMUSG00000105935) | 1.79 | 0.0454 | | | | |
| Gm43667 (ENSMUSG00000105135) | -2.13 | 0.0000 | | | | |
| Gm43689 (ENSMUSG00000104675) | | | -2.03 | 0.0122 | | |
| Gm44371 (ENSMUSG00000107753) | -1.53 | 0.0046 | | | | |
| Gm44681 (ENSMUSG00000109256) | | | -1.31 | 0.0491 | | |
| Gm44812 (ENSMUSG00000109021) | -1.02 | 0.0103 | | | | |
| Gm44896 (ENSMUSG00000108581) | | | -2.12 | 0.0112 | | |
| Gm45201 (ENSMUSG00000109031) | | | -1.89 | 0.0300 | | |
| Gm45206 (ENSMUSG00000108389) | | | -1.14 | 0.0000 | -1.04 | 0.0000 |
| Gm45231 (ENSMUSG00000108888) | | | -1.05 | 0.0033 | | |
| Gm45259 (ENSMUSG00000110050) | | | -1.60 | 0.0446 | -1.24 | 0.0404 |
| Gm45324 (ENSMUSG00000110147) | -1.35 | 0.0090 | | | | |
| Gm45623 (ENSMUSG00000110086) | -1.41 | 0.0403 | | | | |
| Gm45644 (ENSMUSG00000109636) | | | -1.01 | 0.0272 | | |
| Gm45698 (ENSMUSG00000109428) | -1.43 | 0.0496 | | | | |
| Gm45753 (ENSMUSG00000031640) | -1.70 | 0.0045 | | | | |
| Gm46519 (ENSMUSG00000115969) | | | 1.28 | 0.0025 | 1.17 | 0.0098 |
| Gm47270 (ENSMUSG00000111722) | 1.84 | 0.0026 | | | | |
| Gm47467 (ENSMUSG00000113019) | | | 1.94 | 0.0057 | | |
| Gm47621 (ENSMUSG00000112576) | -1.47 | 0.0492 | | | | |
| Gm4779 (ENSMUSG00000045010) | -2.13 | 0.0204 | -2.29 | 0.0090 | -2.06 | 0.0313 |
| Gm47987 (ENSMUSG00000113698) | 1.89 | 0.0399 | | | | |
| Gm4815 (ENSMUSG00000114733) | | | | | 2.03 | 0.0168 |
| Gm48420 (ENSMUSG00000113070) | -1.31 | 0.0419 | | | | |
| Gm48737 (ENSMUSG00000111107) | | | | | -1.58 | 0.0006 |
| Gm48985 (ENSMUSG00000115223) | 2.47 | 0.0244 | | | | |
| Gm5422 (ENSMUSG00000039684) | | | 1.68 | 0.0040 | | |
| Gm5544 (ENSMUSG00000074388) | 1.06 | 0.0000 | | | | |
| Gm5829 (ENSMUSG00000101834) | | | | | 1.98 | 0.0098 |
| Gm5860 (ENSMUSG00000085022) | | | 1.68 | 0.0280 | | |
| Gm6089 (ENSMUSG00000105422) | -1.32 | 0.0146 | -1.55 | 0.0056 | -1.08 | 0.0432 |
| Gm6283 (ENSMUSG00000098057) | -1.96 | 0.0067 | -1.79 | 0.0108 | -1.95 | 0.0186 |
| Gm7285 (ENSMUSG00000105408) | 2.19 | 0.0042 | | | | |
| Gm8615 (ENSMUSG00000024437) | | | -2.29 | 0.0323 | | |
| Gpr156 (ENSMUSG00000046961) | | | 1.54 | 0.0366 | | |
| Gpsm2 (ENSMUSG00000027883) | | | 1.31 | 0.0019 | | |
| Gsdme (ENSMUSG00000029821) | 1.14 | 0.0025 | | | | |
| Hba-a1 (ENSMUSG00000069919) | | | 2.34 | 0.0002 | | |
| Hba-a2 (ENSMUSG00000069917) | | | 2.29 | 0.0004 | | |
| Hbb-bs (ENSMUSG00000052305) | | | 2.23 | 0.0007 | | |
| Hbb-bt (ENSMUSG00000073940) | | | 2.31 | 0.0004 | | |
| Hbq1b (ENSMUSG00000073063) | | | 1.56 | 0.0086 | | |
| Hip1 (ENSMUSG00000039959) | -1.46 | 0.0039 | | | | |
| Hist1h2ac (ENSMUSG00000069270) | -1.75 | 0.0314 | | | | |
| Hyal1 (ENSMUSG00000010051) | -1.19 | 0.0392 | | | | |
| Hykk (ENSMUSG00000035878) | -1.58 | 0.0398 | | | | |
| Ido2 (ENSMUSG00000031549) | 1.73 | 0.0450 | | | | |
| Ifit3 (ENSMUSG00000074896) | | | 1.85 | 0.0124 | | |
| Il17rb (ENSMUSG00000015966) | | | 1.87 | 0.0130 | | |
| Irf9 (ENSMUSG00000002325) | | | 1.61 | 0.0002 | | |
| Isg20 (ENSMUSG00000039236) | | | 2.00 | 0.0491 | | |
| Kcnmb1 (ENSMUSG00000020155) | -1.97 | 0.0000 | | | | |
| Kdelr3 (ENSMUSG00000010830) | -1.38 | 0.0102 | | | | |
| Kdm5d (ENSMUSG00000056673) | | | | | -1.38 | 0.0334 |
| Kirrel (ENSMUSG00000041734) | | | | | -1.11 | 0.0411 |
| Ldb3 (ENSMUSG00000021798) | 2.08 | 0.0001 | 2.16 | 0.0000 | 1.94 | 0.0012 |
| Lgals3 (ENSMUSG00000050335) | | | 2.33 | 0.0000 | | |
| Lhpp (ENSMUSG00000030946) | 1.08 | 0.0105 | | | | |
| Lyz2 (ENSMUSG00000069516) | | | 1.95 | 0.0204 | | |
| Mbd6 (ENSMUSG00000025409) | -1.71 | 0.0491 | | | | |
| Mbtd1 (ENSMUSG00000059474) | -1.16 | 0.0382 | | | -1.41 | 0.0090 |
| Mccc2 (ENSMUSG00000021646) | 1.19 | 0.0024 | | | | |
| Mgp (ENSMUSG00000030218) | | | 2.07 | 0.0044 | | |
| Mid1 (ENSMUSG00000035299) | -1.95 | 0.0250 | | | | |
| Mindy4b-ps (ENSMUSG00000101860) | 1.22 | 0.0197 | | | | |
| Mir5113 (ENSMUSG00000092868) | -1.74 | 0.0398 | | | | |
| Mir7654 (ENSMUSG00000098917) | -1.27 | 0.0260 | | | | |
| Mki67 (ENSMUSG00000031004) | -1.70 | 0.0291 | | | | |
| Mks1 (ENSMUSG00000034121) | 1.76 | 0.0119 | | | | |
| Mroh2a (ENSMUSG00000079429) | -1.17 | 0.0163 | | | | |
| Mtfr1 (ENSMUSG00000027601) | | | 1.11 | 0.0428 | | |
| Muc3a (ENSMUSG00000094840) | -2.14 | 0.0019 | | | | |
| Mus81 (ENSMUSG00000024906) | 1.59 | 0.0000 | | | | |
| Myb (ENSMUSG00000019982) | | | 1.03 | 0.0298 | | |
| Ngp (ENSMUSG00000032484) | | | 1.06 | 0.0000 | | |
| Nlrp5-ps (ENSMUSG00000041596) | 1.33 | 0.0398 | | | | |
| Nmbr (ENSMUSG00000019865) | | | 1.29 | 0.0240 | | |
| Nnat (ENSMUSG00000067786) | | | 1.22 | 0.0240 | | |
| Nxph2 (ENSMUSG00000069132) | | | | | 1.05 | 0.0005 |
| Onecut2 (ENSMUSG00000045991) | -2.42 | 0.0000 | | | | |
| Ovol1 (ENSMUSG00000024922) | -1.45 | 0.0095 | | | | |
| Pacc1 (ENSMUSG00000026627) | | | 1.04 | 0.0446 | | |
| Pcgf2 (ENSMUSG00000018537) | | | -1.38 | 0.0193 | | |
| Pcolce2 (ENSMUSG00000015354) | -1.77 | 0.0190 | | | | |
| Pde3a (ENSMUSG00000041741) | -2.09 | 0.0007 | | | | |
| Pdgfd (ENSMUSG00000032006) | | | 1.80 | 0.0440 | | |
| Pgm3 (ENSMUSG00000056131) | 1.44 | 0.0000 | | | | |
| Pi15 (ENSMUSG00000067780) | | | | | -1.24 | 0.0092 |
| Pidd1 (ENSMUSG00000025507) | -1.51 | 0.0041 | | | | |
| Plbd1 (ENSMUSG00000030214) | | | 1.35 | 0.0087 | | |
| Plscr2 (ENSMUSG00000032372) | | | 2.06 | 0.0215 | | |
| Ppat (ENSMUSG00000029246) | 1.11 | 0.0490 | | | | |
| Prg4 (ENSMUSG00000006014) | | | 1.74 | 0.0072 | | |
| Prps1l3 (ENSMUSG00000079104) | | | 1.14 | 0.0243 | | |
| Prss35 (ENSMUSG00000033491) | -1.06 | 0.0000 | | | | |
| Rab37 (ENSMUSG00000020732) | -1.47 | 0.0000 | | | | |
| Raly (ENSMUSG00000027593) | 1.28 | 0.0044 | | | | |
| Rars2 (ENSMUSG00000028292) | 1.04 | 0.0031 | | | | |
| Rcl1 (ENSMUSG00000024785) | 1.06 | 0.0386 | | | | |
| Rilp (ENSMUSG00000038195) | | | 1.58 | 0.0478 | | |
| Rnase1 (ENSMUSG00000035896) | | | | | -1.01 | 0.0045 |
| Rnf135 (ENSMUSG00000020707) | 1.37 | 0.0454 | | | | |
| Rnf138rt1 (ENSMUSG00000083695) | 1.26 | 0.0161 | | | | |
| Rraga (ENSMUSG00000070934) | 1.25 | 0.0009 | | | | |
| Scube3 (ENSMUSG00000038677) | -1.49 | 0.0074 | | | | |
| Sema5a (ENSMUSG00000022231) | -1.93 | 0.0119 | | | | |
| Setdb2 (ENSMUSG00000071350) | | | 1.80 | 0.0045 | | |
| Slc22a8 (ENSMUSG00000063796) | | | 1.36 | 0.0493 | | |
| Smarcad1 (ENSMUSG00000029920) | 1.26 | 0.0183 | | | | |
| Snora52 (ENSMUSG00000064666) | 1.52 | 0.0244 | | | | |
| Sp140 (ENSMUSG00000070031) | | | 2.68 | 0.0002 | | |
| Spx (ENSMUSG00000071112) | 1.30 | 0.0180 | 2.54 | 0.0000 | 1.64 | 0.0063 |
| Ston1 (ENSMUSG00000033855) | -2.24 | 0.0424 | | | | |
| Taf8 (ENSMUSG00000023980) | | | | | 1.19 | 0.0133 |
| Thbd (ENSMUSG00000074743) | | | | | -1.67 | 0.0294 |
| Tifa (ENSMUSG00000046688) | | | 1.47 | 0.0270 | | |
| Tll1 (ENSMUSG00000053626) | 2.30 | 0.0001 | 1.32 | 0.0163 | | |
| Tlr1 (ENSMUSG00000044827) | -1.08 | 0.0386 | | | | |
| Tlr13 (ENSMUSG00000033777) | | | 1.38 | 0.0037 | | |
| Tmed9 (ENSMUSG00000058569) | 1.38 | 0.0000 | | | | |
| Tmem150cos (ENSMUSG00000086259) | | | -1.22 | 0.0262 | | |
| Tns3 (ENSMUSG00000020422) | -1.48 | 0.0488 | | | | |
| Tpm4 (ENSMUSG00000031799) | | | 1.08 | 0.0090 | | |
| Trim80 (ENSMUSG00000070332) | 1.03 | 0.0000 | | | | |
| Ttc12 (ENSMUSG00000040219) | | | 1.33 | 0.0007 | | |
| Ttc38 (ENSMUSG00000035944) | -1.74 | 0.0496 | | | | |
| Ttll4 (ENSMUSG00000033257) | -2.11 | 0.0031 | | | | |
| Ube2l6 (ENSMUSG00000027078) | | | 1.57 | 0.0301 | | |
| Vmn1r4 (ENSMUSG00000071893) | | | | | 1.27 | 0.0191 |
| Wdfy2 (ENSMUSG00000014547) | -1.71 | 0.0341 | | | | |
| Wdr93 (ENSMUSG00000039099) | | | -1.18 | 0.0256 | | |
| Wfdc17 (ENSMUSG00000069792) | | | 2.36 | 0.0129 | | |
| Wnt11 (ENSMUSG00000015957) | | | 1.08 | 0.0136 | | |
| Wnt16 (ENSMUSG00000029671) | 1.98 | 0.0000 | | | | |
| Wnt2 (ENSMUSG00000010797) | -1.41 | 0.0000 | | | | |
| Zbtb32 (ENSMUSG00000006310) | | | 1.52 | 0.0489 | 1.84 | 0.0288 |
| Zc3h7b (ENSMUSG00000022390) | -1.45 | 0.0202 | | | | |
| Zfp503 (ENSMUSG00000039081) | 1.67 | 0.0179 | | | | |
| Zfp947 (ENSMUSG00000063383) | | | 1.59 | 0.0110 | | |
| Zscan26 (ENSMUSG00000022228) | -1.09 | 0.0212 | | | | |
