## Supplementary material for "Reconceptualising resilience within a translational framework is supported by unique and brain-region specific transcriptional signatures in mice": Table S1 Ventral Hippocampus

| | Indiscriminate-avoiders | | Discriminating-avoiders | |
|---|---|---|---|---|
| **MGI symbol (Ensembl gene ID)** | *log2-fold-change* | *adjusted p-value* | *log2-fold-change* | *adjusted p-value* |
| Ace (ENSMUSG00000020681) | | | -2.78 | 0 |
| Adgrl2 (ENSMUSG00000028184) | | | -2.14 | 0 |
| Chrm5 (ENSMUSG00000074939) | | | -1.78 | 0.0380 |
| Cldn9 (ENSMUSG00000066720) | | | -1.28 | 0.0260 |
| Gipr (ENSMUSG00000030406) | | | -1.44 | 0.0331 |
| Gm11651 (ENSMUSG00000085297) | | | -2.65 | 0.0040 |
| Gm14133 (ENSMUSG00000087029) | | | 1.89 | 0.0392 |
| Gm42865 (ENSMUSG00000104807) | | | 1.19 | 0.0034 |
| Hba-a1 (ENSMUSG00000069919) | 1.64 | 0.0351 | | |
| Hba-a2 (ENSMUSG00000069917) | 1.62 | 0.0435 | | |
| Hbb-bs (ENSMUSG00000052305) | 1.77 | 0.0125 | | |
| Hbb-bt (ENSMUSG00000073940) | 1.82 | 0.0107 | | |
| Hcrt (ENSMUSG00000045471) | 4.11 | 3.2701E-07 | | |
| Hs6st2 (ENSMUSG00000062184) | | | -1.38 | 0.0281 |
| Kcnt1 (ENSMUSG00000058740) | | | -1.73 | 0.0125 |
| Nacad (ENSMUSG00000041073) | | | -1.64 | 0.0083 |
| Nckap5l (ENSMUSG00000023009) | | | -1.56 | 0.0419 |
| Pdzd7 (ENSMUSG00000074818) | | | -1.49 | 0.0098 |
| Phc1 (ENSMUSG00000040669) | | | -1.23 | 0.0260 |
| Slc22a23 (ENSMUSG00000038267) | | | -1.58 | 0.0260 |
| Tbc1d9 (ENSMUSG00000031709) | | | -1.44 | 0.0158 |
| Tctn2 (ENSMUSG00000029386) | | | -1.14 | 0.0471 |
